# SF3B1 Phosphorylation Prompts U2AF2 Dissociation for Widespread Control of pre-mRNA Splicing

**DOI:** 10.64898/2026.01.19.700466

**Authors:** Christopher L. Kirchhoff, Hannah R. Powell, Justin W. Galardi, Sarah Loerch, Mary J. Pulvino, Jermaine L. Jenkins, Paul L. Boutz, Clara L. Kielkopf

## Abstract

Reversible pre-mRNA splicing factor phosphorylation is a well-documented feature of spliceosome assembly, activation, and disassembly, yet its functional and mechanistic roles are still emerging. SF3B1, a core spliceosome subunit, is extensively phosphorylated before the first catalytic step of pre-mRNA splicing. Intriguingly, most SF3B1 phosphorylation sites surround its binding sites for U2AF2, an early-stage pre-mRNA splicing factor. Here, we discovered that SF3B1 phosphorylation significantly decreases its association with U2AF2. We determined two crystal structures revealing electrostatic repulsion between an acidic U2AF2 α-helix and the negatively-charged phosphoryl group of SF3B1. Variants with amino acid substitutions that prevent or mimic SF3B1 phosphorylation perturbed thousands of splice sites, primarily those marked by uridine-rich splice site signals recognized by U2AF2. Collectively, our findings demonstrate a widespread but previously unrecognized role for SF3B1 phosphorylation as a gateway for U2AF2 dissociation so that pre-mRNA splicing can proceed.

---

Pre-messenger RNA splicing, a fundamental process in eukaryotic gene expression, accurately removes introns and ligates exons to form mature mRNA. This intricate process relies on the stepwise assembly of five small nuclear ribonucleoprotein particles (snRNPs) in the spliceosome and is aided by numerous non-snRNP proteins.^1^ To safeguard splicing outcomes, splicing factors are tightly regulated by various post-translational modifications, particularly reversible protein phosphorylation.^2^ Indeed, phosphorylation has been detected for at least one-third of spliceosome subunits, and their phosphorylation status changes across stages of spliceosome assembly, catalytic activation, and disassembly.^3^ Despite its widespread occurrence, the functional and mechanistic consequences of dynamic spliceosome phosphorylations during pre-mRNA splicing remain a major gap in our understanding of gene regulation.

A key player among phosphorylated pre-mRNA splicing factors is Splicing Factor 3B Subunit 1 (SF3B1), an essential component of the U2 snRNP.^1, 4^ SF3B1 defines the 3’ splice site in the first ATP-dependent step of splicing (A-complex) by binding a polypyrimidine tract (PPT) sequence and facilitating base-pairing between the U2 snRNA and the branch point sequence (BPS) of the intron. SF3B1 is extensively and dynamically phosphorylated during spliceosome assembly. Its phosphorylation levels significantly increase at the B-to-B^ACT^ complex transition (just before the first catalytic step of splicing) and remain high through its dissociation from the B*/C complex (between the first and second catalytic steps).^3, 5–7^ While SF3B1 is a reported substrate for both transcription-associated and cell-cycle cyclin-dependent kinases, including CDK7, CDK9, CDK12/13 and CDK1/CDK2,^8–15^ CDK11 has emerged as a primary kinase responsible for SF3B1 phosphorylation during co-transcriptional splicing.^16–18^ Conversely, phosphatases PP1 and PP2A dephosphorylate SF3B1 at the C complex, and this activity is required for the transition to the second catalytic step.^19^ The progressive and regulated phosphorylation of SF3B1 underscores a critical, but as yet not fully understood, role in spliceosome activation.

Intriguingly, most of the identified phosphorylation sites on SF3B1 are located within or near five U2AF Ligand Motifs (ULMs) (**Fig. 1a**). The SF3B1 ULMs serve as binding sites for proteins containing a U2AF Homology Motif (UHM), a protein-protein interaction domain with an RRM-like topology,^20^ including the U2 snRNP Auxiliary Factor 2 (U2AF2), ^21, 22^ its paralogues RBM39/CAPERα^23^ and PUF60,^24^ as well as SPF45,^25^ which is required for the second catalytic step of pre-mRNA splicing, and Tat-SF1,^26, 27^ a chaperone of the U2 snRNP. The spatial and temporal relationship between SF3B1 phosphorylation and its association with U2AF2 suggests a potential interplay that could influence initial identification and subsequent spliceosome assembly at the 3’ splice site. U2AF2 is a foundational subunit at the 3’ splice site of the early (E)-stage spliceosome.^28^ After SF3B1 is recruited, U2AF2 remains loosely associated with A-stage spliceosomes until its levels decrease in the B-complex. U2AF2 is no longer detected at the activated B^act^ or later stage spliceosomes, although SF3B1 remains an integral part of the spliceosome through the first catalytic step of splicing.^3^

**Fig. 1.**
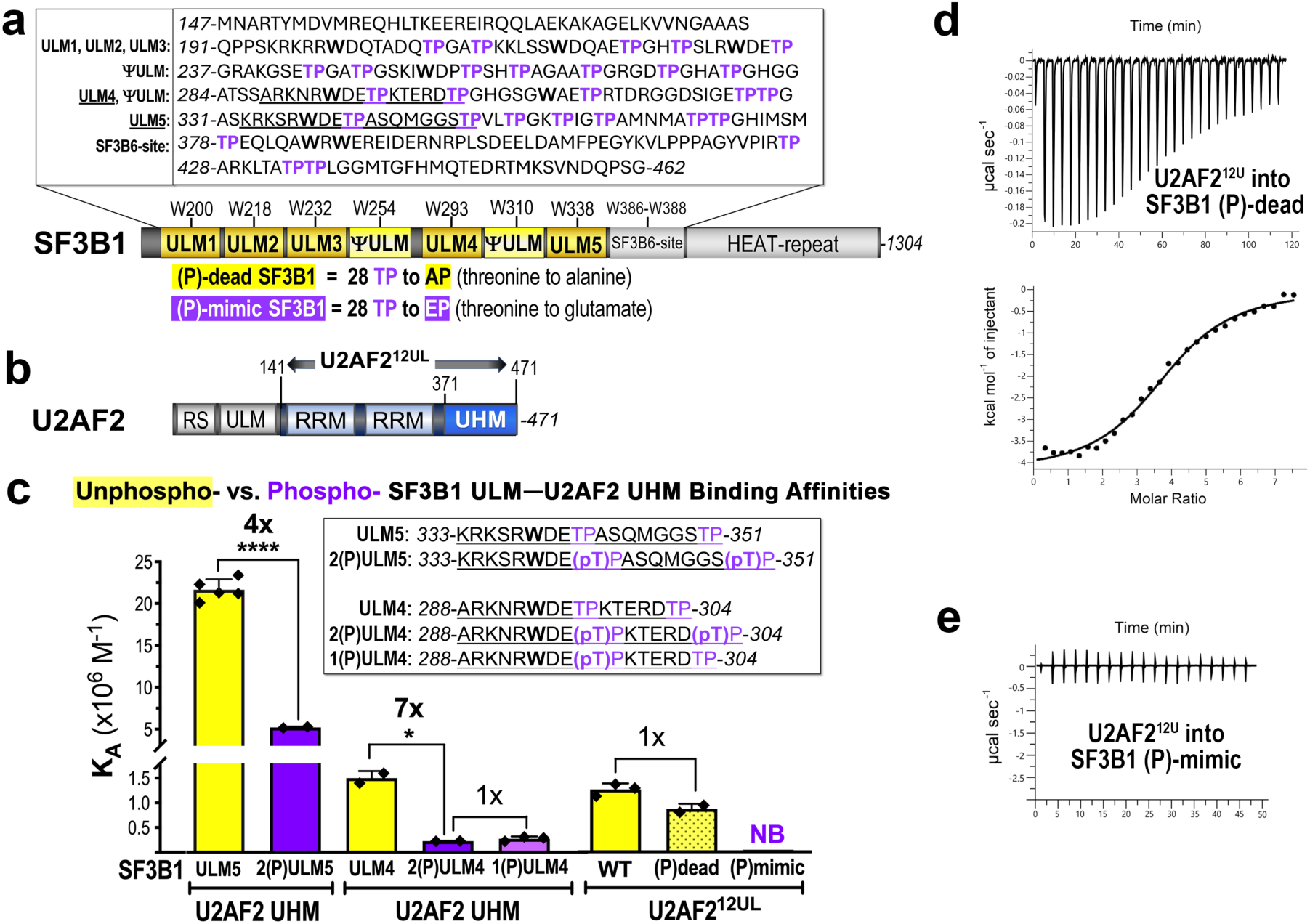
Phosphorylation or (P)-mimics decreased the binding affinity of SF3B1 ULMs for U2AF2. **a** Sites of SF3B1 phosphorylation surround five U2AF Ligand Motifs (ULM) and two ULM-like repeats (ΩULM) (yellow). The SF3B1 variants substitute alanine (TP to AP) or glutamate (TP to EP) for 28 threonine-proline kinase substrates in the ULM region (TP, purple). AlphaFold models of the inset region are shown in **Supplementary** Fig. 2**. b** U2AF2 comprises a C-terminal U2AF Homology Motif (UHM), preceded by two RNA recognition motifs (RRM1 and RRM2) and an N-terminal RS motif. Constructs used for ITC are U2AF2^12L^ (double-headed arrow) and the UHM. **c** Bar graph of the average apparent binding affinities (K_A_) determined by isothermal titration calorimetry with values from separate titrations overlaid (♦) and error bars for standard deviations of these separate experiments. U2AF2 proteins were titrated into unmodified (yellow) or phosphorylated/phospho-mimic (purple) SF3B1 regions or peptides (sequences inset). Welch’s t-test: *, *p* ≤ 0.05; ****, *p* ≤ 0.0001. **d – e** Representative isotherms for the U2AF2^12U<u>L</u>^ titrated into **d** (P)-dead or **e** (P)-mimic SF3B1 variants. The thermodynamic values and other representative isotherms are reported in **Supplementary Table 1** and **Supplementary** Fig. 1.

In the E-stage of spliceosome assembly, U2AF2 initially forms a complex with Splicing Factor-1 (SF1) via a U2AF Homology Motif (UHM),^29^ a protein-protein interaction domain with an RRM-like topology.^20^ In turn, SF1 recognizes the BPS^29^ and two central RNA recognition motifs (RRMs) of U2AF2 identify a uridine-rich PPT signal for the major class of 3’ splice sites.^30^ A heterodimeric partner of U2AF2, called U2AF1, contacts a consensus AG-dinucleotide at the intron-exon junction.^31^ Subsequently, in the first ATP-dependent step of the spliceosome assembly, U2AF2 engages the SF3B1 subunit through a UHM/ULM interface^22^ and recruits the U2 snRNP to the 3’ splice site.^28, 32^ In this A-stage spliceosome, the SF3B1 α-helical repeat (HEAT) region interacts with the PPT,^4^ displacing the U2AF2 RRMs from the pre-mRNA. Concurrently, an N-terminal RS domain of U2AF2 anneals the U2 snRNA with the BPS (**Fig. 1b**).^33^ Crystal structures and biochemical characterizations indicate that an SF3B1 ULM (ULM5) binds the U2AF2 UHM,^22^ thereby facilitating SF1’s exit and the transition to the A-stage spliceosome. Coincident with high levels of SF3B1 phosphorylation at the B^act^ stage, U2AF2 dissociates from the spliceosome.^3^ Despite a plethora of SF3B1-containing structures,^4^ how this exchange of interactions with U2AF2 and the SF3B1 ULM region is orchestrated has not been fully resolved to date.

In this study, we comprehensively investigate the structural and functional consequences of SF3B1 phosphorylation on its interaction with U2AF2 and its broader implications for pre-mRNA splicing. We demonstrate that SF3B1 phosphorylation reduces its association with U2AF2, and we characterize the structural basis for this reduced binding through high-resolution crystal structures. Our transcriptome-wide RNA sequencing shows that variants of SF3B1, which either prevent or mimic phosphorylation, make widespread changes in splicing patterns, particularly affecting splice sites with uridine-rich PPTs recognized by U2AF2. Collectively, our findings establish SF3B1 phosphorylation as a critical and previously unappreciated mechanism governing a U2AF2-dependent gateway for pre-mRNA splicing, thereby providing new insights into the regulatory complexity of this essential cellular process.

## RESULTS

### SF3B1 phosphorylation decreases its binding affinity for U2AF2

To measure the influence of SF3B1 phosphorylation on its binding to U2AF2, we first quantified the binding affinities of recombinant proteins using isothermal titration calorimetry (ITC) (**Fig. 1c-e**, **Supplementary Fig. 1, Supplementary Table 1**). We titrated the U2AF2 UHM, which is the known SF3B1-interaction domain,^22^ into SF3B1 ULM peptides with defined phosphorylation sites. Phosphorylation of the cognate SF3B1 ULM (ULM5, residues 333-351) reduced its apparent equilibrium binding affinity (*K_A_*) for the U2AF2 UHM by four-fold, whereas phosphorylation of a less preferred site (ULM4, residues 288-304) reduced the *K_A_* by seven-fold. Phosphorylation of a single TP motif proximal to a signature ULM tryptophan imparted a similar penalty as dual phosphorylation of the two TP motifs in the ULM4 peptide.

Given the extensive phosphorylation of TP motifs clustered at the SF3B1 ULM region in the B^act^ spliceosome,^3, 5^ we investigated whether phosphorylation of all TP motifs in the N-terminal SF3B1 region comprising the ULMs (residues 147-462) would exacerbate the U2AF2 binding penalty. To generate a “phospho (P)-mimic” SF3B1, we replaced all 28 threonines in the TP motifs of this SF3B1 region with glutamate residues, which are known to mimic the charge and size of phosphoryl-threonines (e.g., ^34–36^). We were not concerned that amino acid substitutions would disrupt the protein fold since our previous circular dichroism spectra indicate that the SF3B1 ULMs are an intrinsically disordered region (IDR) without detectable secondary structure.^21^ Moreover, AlphaFold models^37^ of the unmodified, substituted variants, or fully phosphorylated states lacked regular structure for the ULM region (**Supplementary Fig. 2**). We titrated the (P)-mimic SF3B1 region with a lengthened U2AF2 construct (U2AF2^12UL^, residues 141 to the C-terminus), only lacking N-terminal, aggregation-prone motifs. U2AF2^12UL^ binds the intact SF3B1 ULM-region with slightly lower *K_A_* than the preferred ULM5 site,^21, 22^ most likely because the fitted *K_A_* is a mixture of binding affinities for five different SF3B1 ULMs. Remarkably, the (P)-mimic SF3B1 variant no longer bound to U2AF2^12UL^ by ITC (**Fig. 1c**, **e**), indicating at least 25-fold affinity loss considering limits for detection.

To confirm that lack of U2AF2 binding by the (P)-mimic protein was not due to removal of specific threonine interactions, we constructed a “(P)-dead” variant, substituting the threonines in the clustered TP-motifs of the ULM-containing region (residues 147-462) with alanines that could not be phosphorylated (**Fig. 1a**). By ITC, the (P)-dead SF3B1 region bound to U2AF2^12UL^ with similar affinity as the WT SF3B1 protein (**Fig. 1c-d**, **Supplementary Table 1**), indicating that the U2AF2 binding was insensitive to neutral perturbations of the SF3B1 threonines. Altogether, establishment of these (P)-dead and (P)-mimic SF3B1 variants equipped us with tools to examine the functional consequences of SF3B1 phosphorylation for its association with U2AF2 and influence on splicing in human cells.

### SF3B1 phosphorylation selectively decreases its association with U2AF2 in cell lysates

We next asked whether phosphorylation of full length SF3B1 would influence its association with U2AF2 in human cell extracts. Immunoblots for SF3B1 phosphorylation, including specific phosphoryl (P)T211 or (P)T313 sites (near ULM1/ULM2 or ULM4/ULM5, respectively) confirmed that wild-type (WT) FLAG-tagged SF3B1 is phosphorylated in our lysis conditions (**Fig. 2a-b, Supplementary Fig. 3**). Treatment with lambda protein phosphatase (^α^PPase) abolished detectable phosphorylation of the T211 or T313 sites (**Fig. 2a-b**). Consistent with broad phosphatase activity and phosphorylation sites in the U2AF2 RS domain, an HA-tagged band with altered migration appeared after incubation with ^α^PPase, suggesting partial dephosphorylation of ^HA^U2AF2. However, this species was not apparent in the presence of the SF3B1 variants (e.g., **Fig. 2d-e**), and unphosphorylated U2AF2 is known to support pre-mRNA splicing in other contexts.^38^ As expected, phosphorylation of the mutant T211A/T211E or T313A/T311E sites was not detected for the (P)-dead and (P)-mimic S3B1 variants (**Fig. 2a-b**). These variants reduced overall TP phosphorylation of the SF3B1 protein to background levels in immunoprecipitates of human cell extracts, confirming that essentially all detected phosphorylated TP sites of SF3B1 fall within the ULM region (**Supplementary Fig. 3**).

**Fig. 2.**
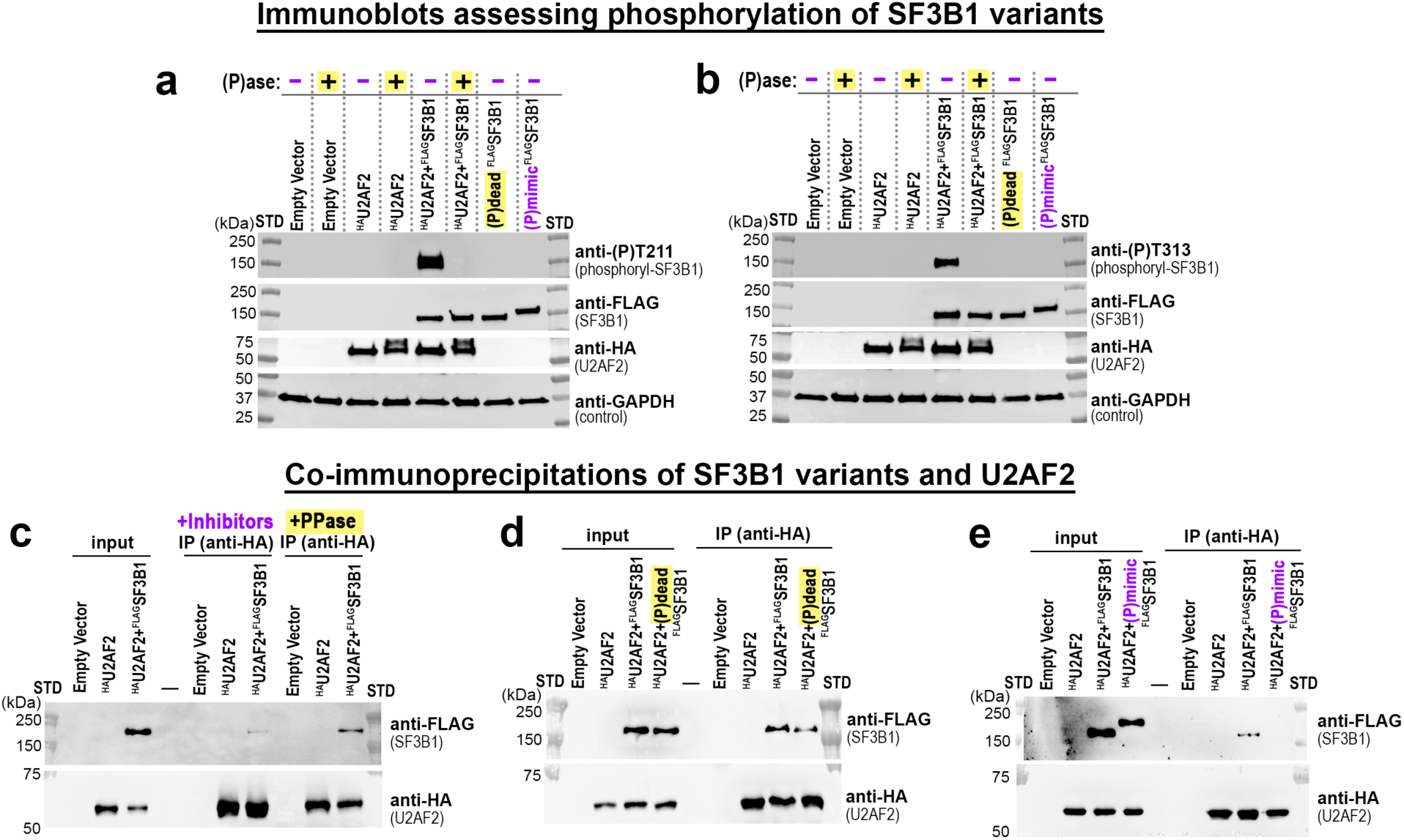
Phosphorylation of the SF3B1 ULM region decreases association with U2AF2. **a – b** Immunoblots confirming that phosphatase treatment dephosphorylated SF3B1 and the (P)-dead or (P)-mimic variants were not detectably phosphorylated at **a** T211 or **b** T313 sites. Total TP phosphorylation is assessed in **Supplementary Fig. 3**. **c – e** Co-immunoprecipitations of HA-tagged U2AF2 with either **c** PPase-treated, **d** (P)-dead, or **e** (P)-mimic Flag-tagged SF3B1 transiently expressed in HEK 293T cells. STD, protein molecular weight markers.

The WT SF3B1 protein co-immunoprecipitated (co-IP) with U2AF2 from extracts of transiently transfected HEK 293T cells (**Fig. 2c**), as observed previously.^22^ This WT SF3B1 retention of U2AF2 increased when ^α^PPase was included in the co-IPs. (P)-dead SF3B1 showed similar association with U2AF2 as the WT SF3B1 counterpart (**Fig. 2d**), whereas in agreement with its low binding affinity, the (P)-mimic SF3B1 variant did not detectably immunoprecipitate with U2AF2 (**Fig. 2e**). To determine if these effects were specific to U2AF2, we compared co-IPs of the (P)-mimic SF3B1 variant with other UHM-domain containing factors, TAT-SF1 and RBM17 (SPF45). In contrast to our observations with U2AF2, both TAT-SF1 and RBM17 maintained consistent association with the (P)-mimic SF3B1 variant (Supplementary Fig. X4). These results demonstrate that the clustered TP mutations do not globally disrupt the ULM-containing region, as the associations of TAT-SF1 and RBM17 were markedly less sensitive to these variants than was the association of U2AF2. Collectively, these data suggest that SF3B1 phosphorylation does not function as a universal inhibitor of UHM interactions, but rather serves as a selective regulator to promote U2AF2 dissociation.

### Phosphorylated SF3B1 TP motif is repulsed by an acidic α-helix of U2AF2 UHM

To investigate the structural mechanism for phosphorylation reducing SF3B1 association with U2AF2, we determined crystal structures of two different phosphorylated ULMs bound to U2AF2 (**Fig. 3, Supplementary Table 2,** ULM sequences inset in **Fig. 1c**). We first determined the 1.43 Å resolution structure of the U2AF2 UHM with its preferred ULM5 binding site, for which both TP motifs were phosphorylated. Only the proximal phosphorylation site was evident in the electron density maps of two (P)ULM5 – U2AF2 UHM complexes present in the asymmetric unit (ASU) (**Fig. 3j**). The two crystallographically-independent complexes shared similar conformations with one another and with the structure of the unphosphorylated ULM5 complex (comparing PDB ID 7SN6 chains B/D, which is free of crystal packing with the ULM TP site^22^) (**Fig. 3a-b**). The TP motif remains in a nearly identical location following phosphorylation, placing the negatively-charged phosphoryl group unfavorably close to E405 in an acidic α-helix of the UHM (**Fig. 3e-f**). A conserved lysine (K408) at the C-terminus of the UHM α-helix is positioned to stabilize the location of the ULM phosphoryl group, despite the neighboring, negatively-charged side chains.

**Fig. 3.**
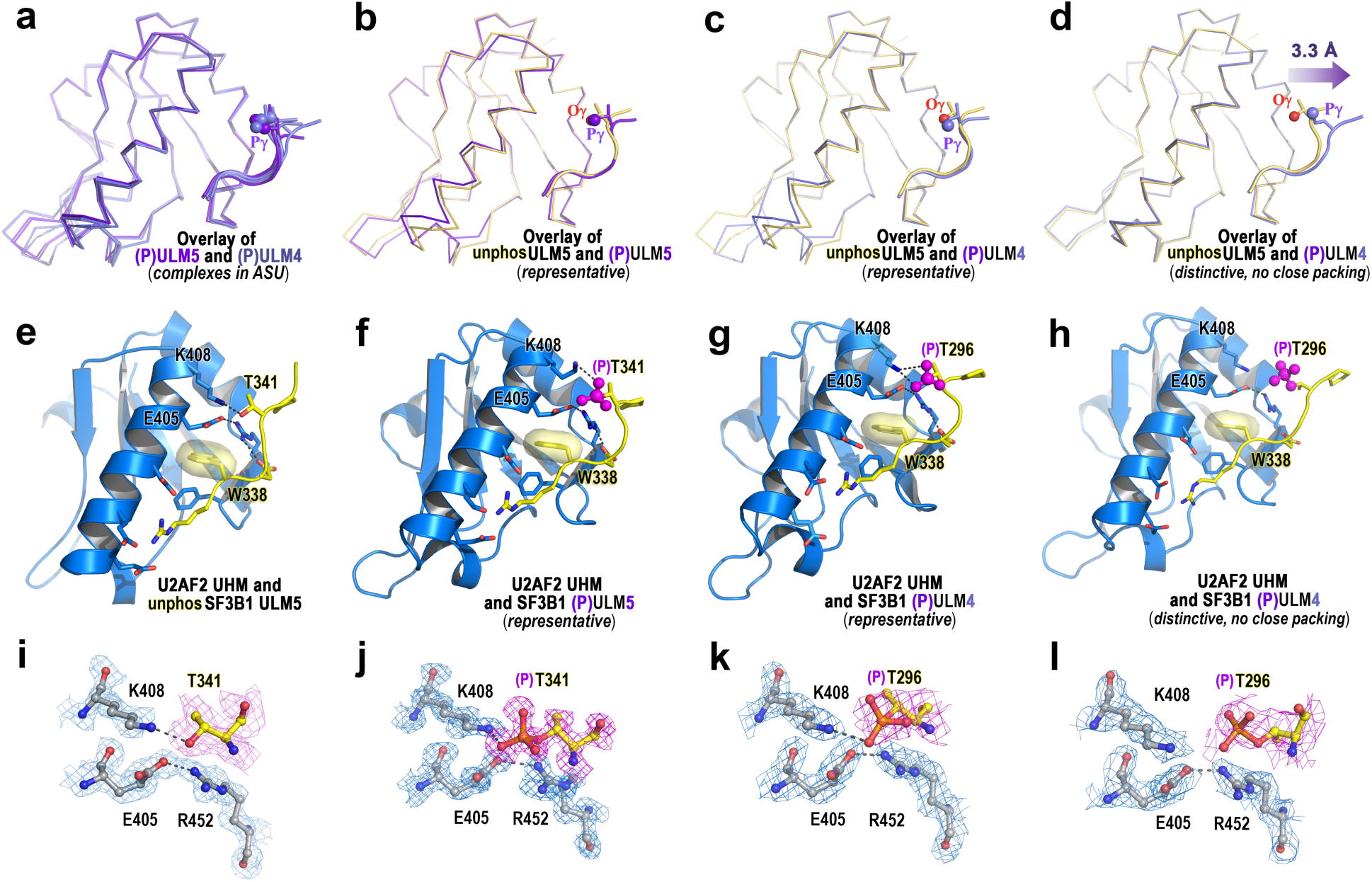
Crystal structures of the U2AF2 UHM bound to phosphorylated SF3B1 ULMs. **a** Superposition of two copies of U2AF2 UHM bound to phosphorylated (P)ULM5 (dark purple) and four copies bound to (P)ULM4 (lilac) in the asymmetric units of the crystal structures. **b** The U2AF2 UHM bound to unphosphorylated ULM5 (PDB ID 7SN6, chains A/C) superimposed with a typical (P)ULM5 complex or **c** (P)ULM4 complex. **d** The (P)ULM4 complex without packing contacts (within 10 Å) shows movement of the phosphorylation site away from the UHM. For **a – d**, spheres mark phosphate atoms or the side chain oxygen (Oψ, red) of unphosphorylated threonine. (**e** – **h**) Overall structures showing (P)T340 interactions. (**i** – **l**) Feature enhanced (bias-reduced) electron density maps of (P)T340 and key neighboring residues for the indicated complexes. Dashed lines connect polar atoms within expected hydrogen bond distance. Crystallographic data and refinement statistics are given in **Supplementary Table 2**.

The trajectory of the bound ULM5 and (P)ULM5 across the U2AF2 UHM is unusual for ULM – UHM complexes, running parallel rather than across the acidic UHM α-helix.^20, 22^ To investigate the effect of varying the ULM sequence on its conformation, we determined the 2.30 Å resolution crystal structure of the U2AF2 UHM with a singly-phosphorylated ULM4 (**Supplementary Table 2**, sequence shown in **Fig. 1c**) This (P)ULM4 had a similar binding affinity as the doubly phosphorylated counterpart (**Supplementary Table 1**). Four crystallographically-independent complexes were present in the ASU of the (P)ULM4 complex. Three of the (P)ULM4 – U2AF2 UHM complexes shared nearly identical conformations with one another and the (P)ULM5-bound structures, with the ULM phosphoryl group interacting with K408 in the acidic UHM α-helix (**Fig. 3a, d**). Although none of the phosphorylated ULMs participated in crystal packing contacts, the phosphoryl group of one of the (P)ULM4 – U2AF2 UHM complexes was especially far from its neighbors. This phosphorylated TP motif had moved 3 Å away from the acidic UHM α-helix compared to the unphosphorylated ULM5 (3.3 Å Oψ – Oψ distance between the superimposed complexes) (**Fig. 3d, h**). The electron density map for this unrestrained (P)TP motif was less defined compared to others (**Fig. 3l**), possibly due to greater mobility. This result indicates that where permitted, the phosphorylated TP motif will dissociate from the negatively-charged UHM surface.

### SF3B1 TP-motifs are critical for SF3B1-sensitive pre-mRNA splicing

Equipped with the (P)-dead and (P)-mimic variants, we were positioned to test the importance of these major SF3B1 phosphorylation sites for its pre-mRNA splicing functions. As an initial assessment, we used reverse transcriptase-polymerase chain reaction (RT-PCR) to assess exon skipping in three well-characterized endogenous transcripts (THYN1, SAT1, RNF10), previously identified as sensitive to SF3B1 levels^22^ (**Fig. 4, Supplementary Fig. 5**). Consistent with prior findings, SF3B1 depletion by siRNA significantly reduced inclusion of cassette exons in these transcripts. Notably, while re-expression of WT SF3B1 from an siRNA-resistant plasmid restored inclusion of these cassette exons, neither the (P)-dead nor the (P)-mimic SF3B1 variants could rescue the altered splicing events.

**Fig. 4.**
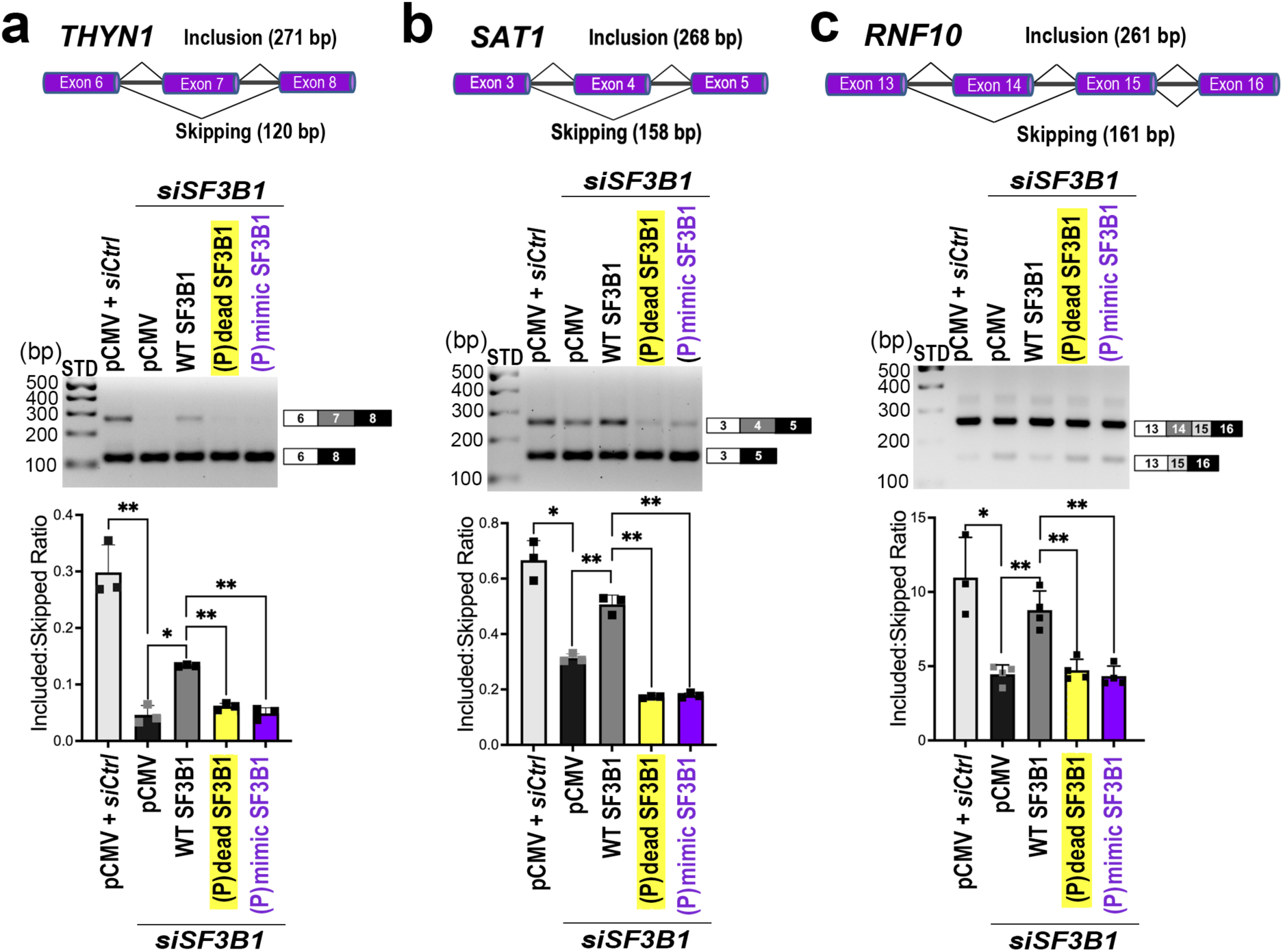
SF3B1 (P)-variants are unable to rescue SF3B1-sensitive pre-mRNA splicing events. RT-PCR of **a** *THYN1*, **b** *SAT1*, or **c** *RNF10* transcripts isolated from human cells treated with either control siRNA (siCtrl, light gray) or SF3B1 siRNA (siSF3B1) and transfected with empty vector (pCMV, black), WT (dark gray), (P)-dead (yellow), or (P)-mimic SF3B1-expressing plasmids (purple). The background-corrected and averaged ratios of exon-included to exon-skipped band intensities from quantification of three separate reactions are shown with error bars for standard deviations and overlays of the individual results (▪) below representative gels of each sample. STD, 100 bp standards. The effect of re-expressing (P)-variant SF3B1 is more severe than SF3B1 depletion for *SAT1* splicing and has no significant difference for *THYN1* or *RNF10* (*p* ≥ 0.25). Welch’s t-test is shown for other comparisons: *, *p* ≤ 0.01; **, *p* ≤ 0.005. Immunoblots and uncropped gel images of all replicates are shown in **Supplementary Fig**. 4.

Building on these targeted results, we next performed transcriptome-wide RNA sequencing (RNA-seq) to globally assess the impact of SF3B1 phosphorylation variants on pre-mRNA splicing. We sequenced RNA samples (RNA-seq) from human HEK 293T cells in which endogenous SF3B1 was depleted and replaced by reexpressed WT, (P)-dead, or (P)-mimic variants (**Fig. 5, Supplementary Fig. 6, Supplementary Data 1-5**). Our depletion strategy compared transfections with two different SF3B1-directed siRNAs after 48 hours. This timeframe efficiently reduced SF3B1 levels (*SF3B1* mRNA knockdown efficiencies 79% for siSF3B1a and 74% for siSF3B1b) without detectable cell death, as confirmed by the absence of CASP3/PARP1 cleavage in immunoblots (**Supplementary Fig. 6c-f**). SF3B1 depletion resulted in over 4,400 shared differential splicing events, including the *THYN1, SAT1,* and *RNF10* exons (**Supplementary Fig. 6a-b**). Given significant overlap in differential splicing between the two siRNAs, we focused on the more efficient siRNA for subsequent re-expression experiments. In agreement with the RT-PCR results, WT SF3B1 re-expression significantly restored inclusion of the *THYN1, SAT1,* or *RNF10* cassette exons, whereas these exons remained skipped in the (P)-dead or (P)-mimic SF3B1 samples (**Fig. 5a-b**).

**Fig. 5.**
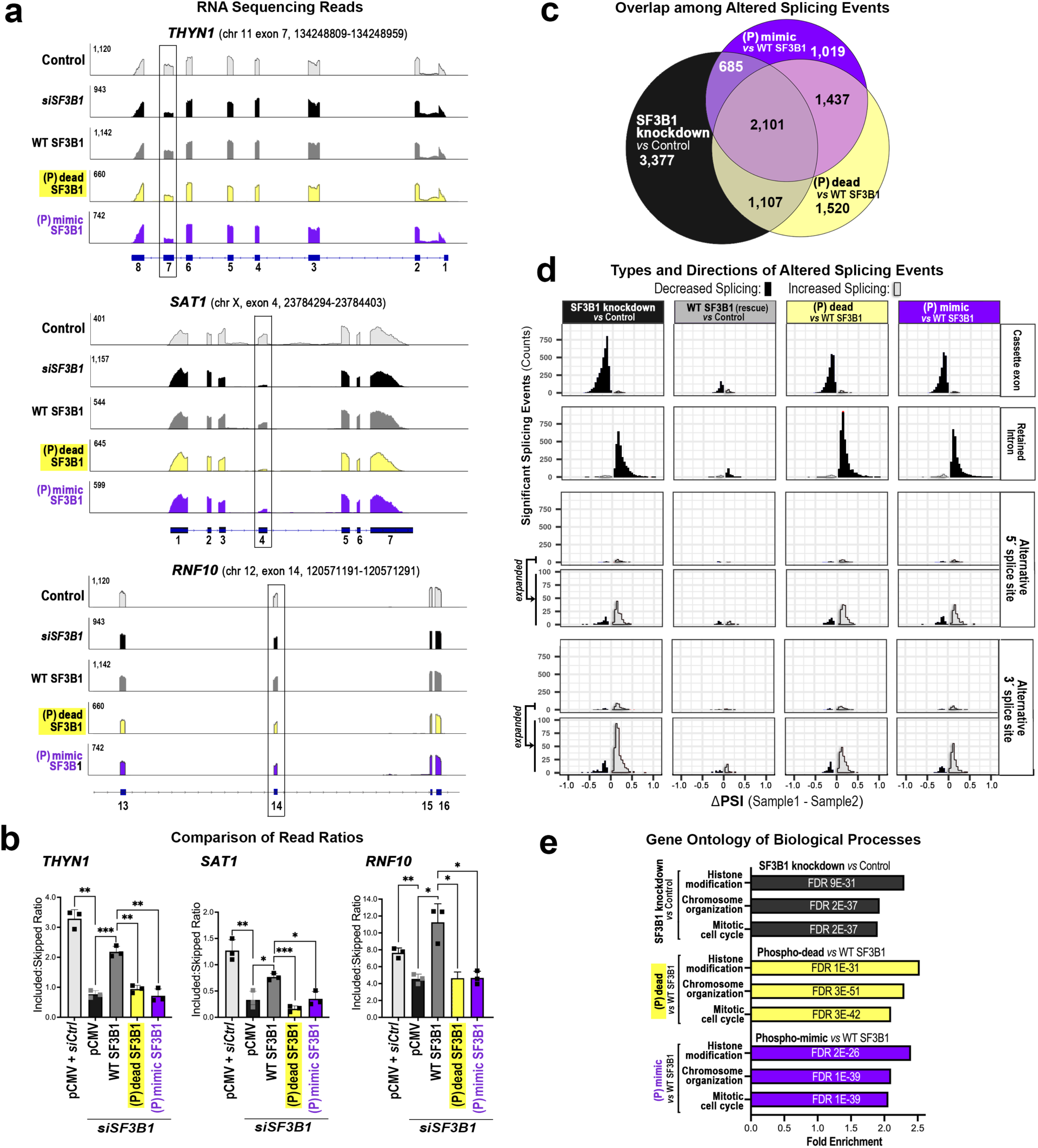
Phospho-dead or phospho-mimic SF3B1 variants have transcriptome-wide effects on pre-mRNA splicing including those due to loss of SF3B1. **a** Representative RNA sequencing reads and **b** quantified read counts of representative SF3B1-sensitive transcripts (*SAT1*, *THYN1*, or *RNF10*). The quantified read counts from three separate transfections per sample are individually overlaid on bar graphs of the average values with error bars for the standard deviations. Welch’s unpaired t-tests: n.s., not significant; *, *p* ≤ 0.05; **, *p* ≤ 0.005; ***, *p* ≤ 0.001. **c** Euler diagram of overlap of significant splicing differences among SF3B1 knockdown (black, siSF3B1*vs.* siCtrl) and SF3B1 knockdown with either phospho (P)-dead SF3B1 (yellow, *vs.* WT SF3B1) or phospho-mimic SF3B1 (purple, *vs.* WT SF3B1). Overlapping splice events required the same direction of differential change. The Spearman rank correlations (ρ) of the differentially splicing events are 0.61 for (P)-dead overlap with siSF3B1, 0.62 for (P)-mimic overlap with siSF3B1, and 0.71 for (P)-dead overlap with (P)-mimic, where a ρ value of +1 is perfect positive correlation. **d** Types and directions (increase, white; decrease, black) of statistically significant splicing changes quantified by the difference in “percent spliced in” (ΔPSI, where 0.10 is a 10% change in splicing) between the indicated samples. Rare alternative 5’ and 3’ splicing events are expanded in the lower panels. **e** Gene ontology of differentially spliced transcripts. The false discovery rates (FDR) are inset. Quantified read counts, comparison with a second SF3B1 siRNA, and immunoblots of RNAseq samples are shown in **Supplementary Fig. 5**.

Transcriptome-wide analysis revealed significant overlap in differential splicing patterns among SF3B1 depletion and the (P)-variant re-expression experiments (**Fig. 5c**). Specifically, over 2,000 differential splicing events, with consistent directions of change, were shared across the SF3B1 perturbations, including *(i)* SF3B1 depletion (siSF3B1 vs. control siRNA), *(ii)* re-expression of (P)-dead SF3B1 ((P)-dead/siSF3B1 relative to WT SF3B1/siSF3B1), and *(iii)* re-expression of (P)-mimic SF3B1 ((P)-mimic/siSF3B1 relative to WT SF3B1/siSF3B1). For all three cases, cassette exon skipping and intron retention events were the most common types of altered pre-mRNA splicing (**Fig. 5d**). Furthermore, gene ontology analysis of the affected splice sites demonstrated enrichment of similar functional pathways among the SF3B1-depleted and the (P)-variant re-expression samples, including those related to histone modifications, chromatin organization and cell cycle (**Fig. 5e**).

### SF3B1 (P)-variant-affected PPT sequences support a mechanistic link to U2AF2

Considering the known function of U2AF2 to recognize the PPT, we further explored the mechanistic link between SF3B1 phosphorylation and U2AF2 activity by examining the PPT sequences of affected 3’ splice sites transcriptome-wide (**Fig. 6, Supplementary Table 3**). We specifically examined the uridine contents of the PPTs, since uridine-rich sequences are preferentially bound by U2AF2.^30, 39, 40^ The PPTs were higher in uridine content for the large class of cassette exons exhibiting increased skipping (i.e., decreased splicing of the proximal 3’ splice site) following re-expression of either (P)-dead or (P)-mimic SF3B1, as compared to those with no significant change or decreased skipping in these samples (**Fig. 6a, c**). Conversely, for the much smaller class of cassette exons with increased inclusion following (P)-variant expression, the PPTs were significantly lower in uridine content. By comparison, the differences in PPT content for cassette exons affected in either direction by SF3B1 knockdown were insignificant.

**Fig. 6.**
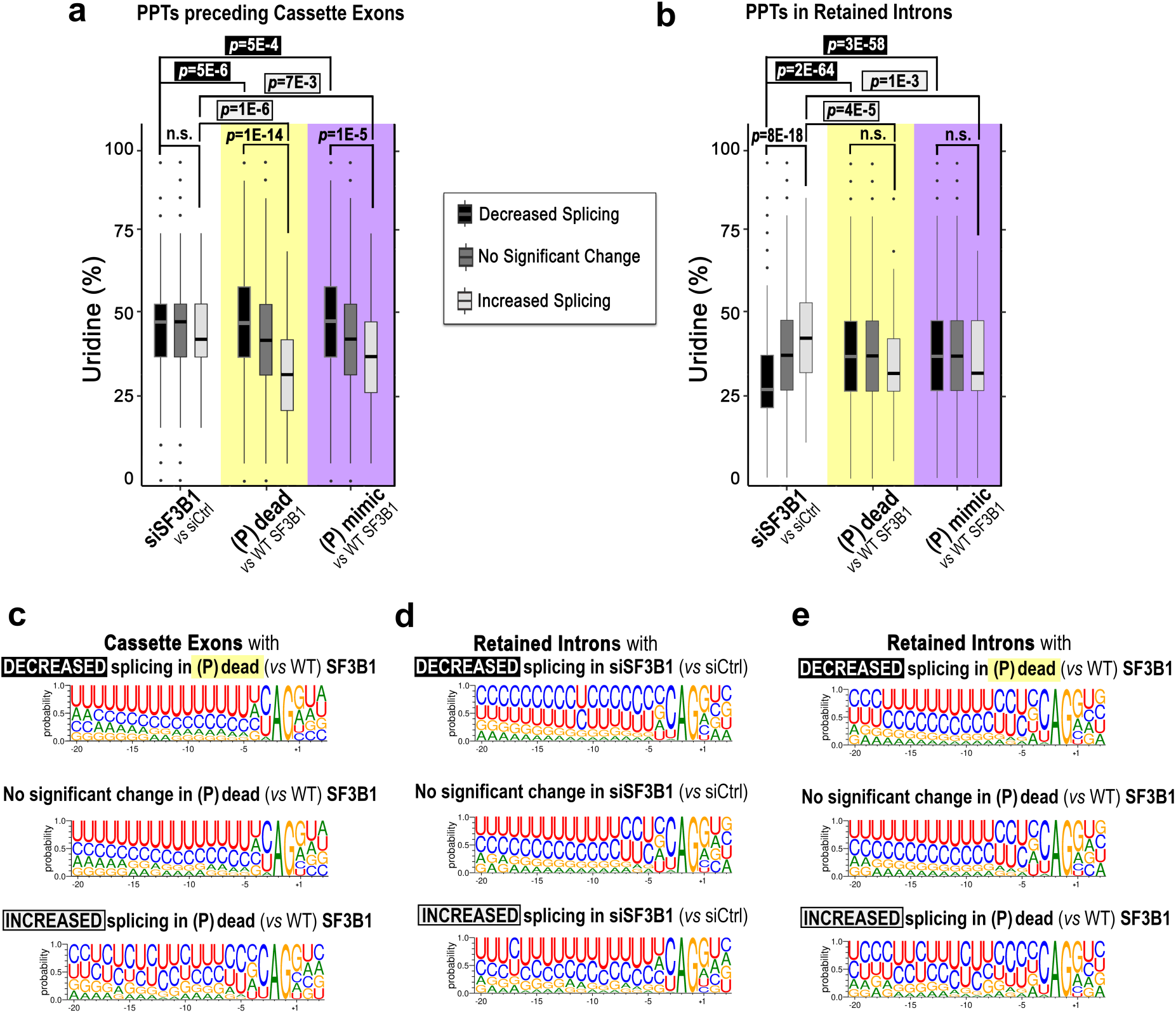
Phospho-variant SF3B1 differently affects 3’ splice sites depending on the uridine contents of the PPTs. **a-b** Box plots compare the uridine percentages for a window (-2 to -20 nucleotides) before 3’ splice sites (SS) located either **a** upstream of cassette exons or **b** in retained introns. Central bars mark the medians (50^th^ percentile), boxes range from 25^th^ percentile to 75^th^ percentile of the data, whiskers extend over 1.5 times the interquartile range, and individual data points mark the outliers. Boxes are shaded by decreased splicing (black), no change (medium gray), or increased splicing (light gray). Columns are colored by sample: white, SF3B1 knockdown (relative to control siRNA); yellow, (P)-dead (relative to WT) SF3B1; violet, (P)-mimic (relative to WT) SF3B1. The *p-*values were calculated by Mann-Whitney U tests. The number of events and *p-*values between samples are detailed in **Supplementary Table 3**. **c-e** Sequence logos of PPTs **c** preceding cassette exons in the (P)-dead (relative to WT) SF3B1, **d** in retained introns of SF3B1 knockdown (relative to control siRNA) or **e** in retained introns of (P)-dead (relative to WT) SF3B1.

The predominant effect on the retained intron class was increased retention (i.e., decreased splicing) across all SF3B1 perturbations, including knockdown and both (P)-variant mutants (**Fig. 5b**). As a whole, the retained intron group had fewer uridines in the PPT compared to the introns flanking cassette exons (**Fig. 6b**). The large class of introns with increased retention following SF3B1 depletion exhibited PPTs with strongly decreased uridine contents, while those of the small group with increased splicing in this sample exhibited higher uridine contents (**Fig. 6b, d**). By contrast, when compared to the SF3B1-depleted samples, the PPTs with increased retention in either the (P)-dead or (P)-mimic SF3B1 samples showed a remarkably significant increase in uridine contents (*p*-values 1.7e-64 and 3.1e-58, **Fig. 6b**,**e**, **Supplementary Table 3**). Although insignificant compared to the large class of introns with increased retention, the median percentages of uridines were lower and the sequence logos relatively degenerate for the small subset of introns with decreased retention in the (P)-variant samples (**Fig. 6b**,**e**). Especially compared to the SF3B1-depleted samples, the different types of splicing events converge on an apparent trend for the (P)-variants to decrease splicing of uridine-rich PPTs characteristic of U2AF2, while increasing splicing of rare PPTs containing lower numbers of uridines.

## DISCUSSION

The ATP-dependence of pre-mRNA splicing has been recognized for more than 40 years, despite thermodynamically favorable transesterification reactions. Historically, the requirement for ATP has been primarily attributed to the rearrangement of ribonucleoproteins by RNA helicases. Protein phosphorylation is now emerging as a second major consumer of ATP for spliceosome activation. Phosphorylation of the key U2 snRNP subunit SF3B1 is a hallmark of active, chromatin-associated spliceosomes,^5^ yet little is known concerning the consequences of SF3B1 phosphorylation for the landscape of pre-mRNA splicing and orchestration of spliceosomal components. We here discover that SF3B1 phosphorylation is a direct regulator of U2AF2 at the entryway to pre-mRNA splicing (**Fig. 7**), contributing to thousands of pre-mRNA splicing events transcriptome-wide.

**Fig. 7.**
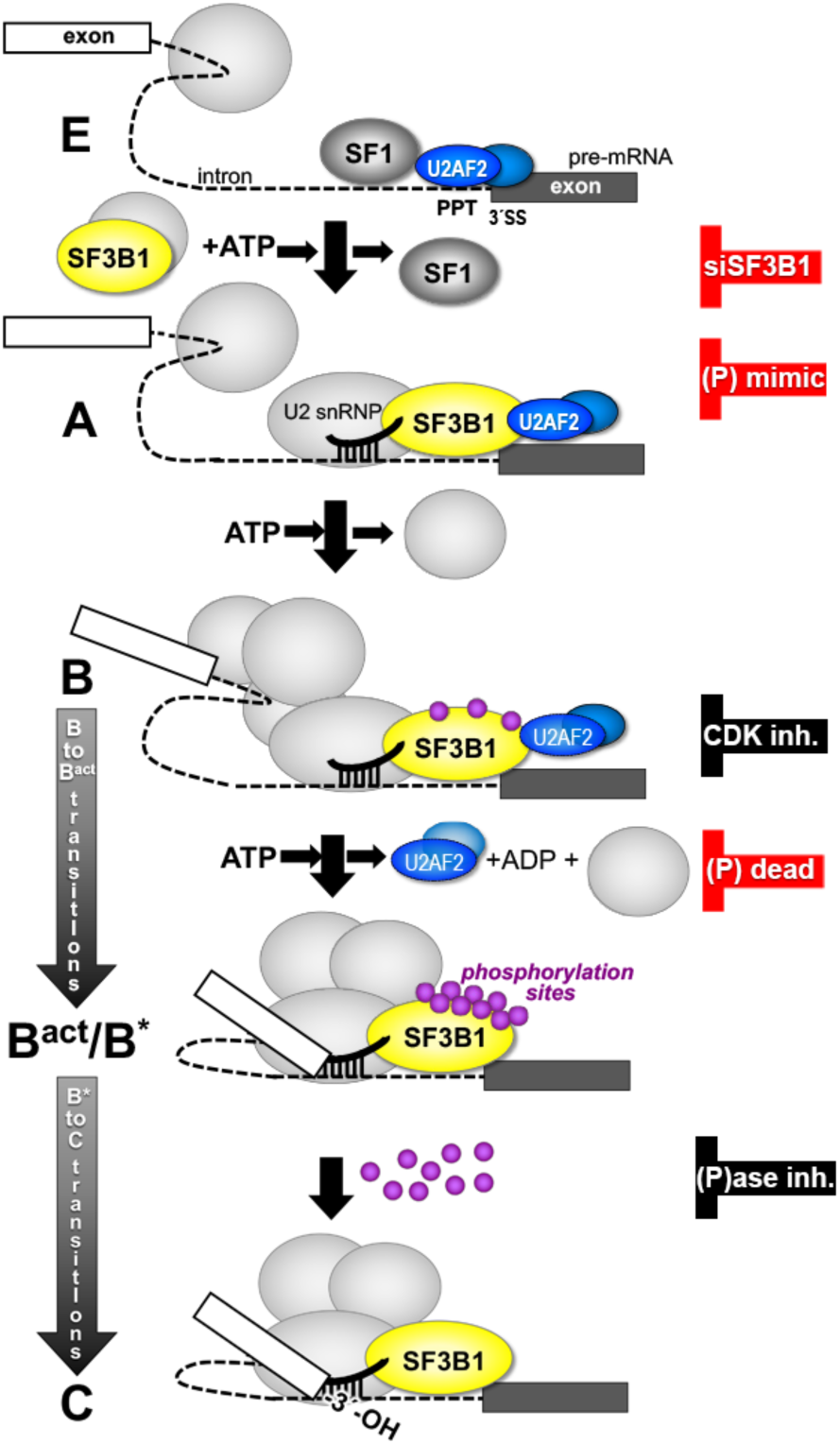
Simplified model for SF3B1 phosphorylation prompting U2AF2 dissociation for spliceosome activation. In the A complex, SF3B1 (yellow ellipse) has low phosphorylation levels and is bound to U2AF2 (blue ellipses) at the PPT of the intron. During activation, SF3B1 becomes highly phosphorylated (purple circles) and U2AF2 dissociates, potentially altering interactions with other splicing factors and RNAs. By the C-complex, SF3B1 is dephosphorylated. Depletion of SF3B1, expression of (P)-dead or (P)-mimic SF3B1 (red symbols), or treatments with kinase or phosphatase inhibitors (inh., black symbols) are expected to block various stages of spliceosome assembly, thereby preventing the catalytic steps of pre-mRNA splicing.

### Phosphorylation of SF3B1 ULMs prompts U2AF2 dissociation

We found that treatments to remove or retain protein phosphorylation or engineered amino acids to prevent or mimic SF3B1 phosphorylation, in turn reciprocally increased or decreased SF3B1 association with U2AF2 in co-immunoprecipitations from cell lysates. While the phosphorylation mimic abolished detectable SF3B1–U2AF2 association, we found that the associations of TAT-SF1 and RBM17 remained largely unaffected (Fig. 2, Supplementary Fig. 4). This differential sensitivity suggests that SF3B1 phosphorylation does not act as a global disruptor of U2 snRNP network. Instead, it appears to function as a selective regulator that displaces U2AF2 while maintaining the recruitment of other UHM-containing factors, perhaps through multivalent binding interfaces or distinct local environments within the disordered ULM region. This selective release may also involve the coordinated recruitment of nuclear phosphatases,^41, 42^ or other spliceosomal subunits to ensure the directional progression of the splicing cycle.^43^

Our high-resolution crystal structures illustrate the phosphorylated TP motif in the ULM positioned near the signature acidic α-helix of the UHM. Comparison among six phosphorylated complexes and the unmodified structure demonstrated the capacity of the U2AF2 UHM to release the phosphorylated SF3B1 group while remaining bound to the ULM. Quantitative ITC measurements using purified protein regions found that TP-phosphorylation decreased the apparent binding affinities of the SF3B1 ULMs for the U2AF2 UHM by four- to seven-fold. Considering its interactions with the acidic UHM α-helix, an additional basic residue in the positively-charged ULM5 tail (K333 compared to A288 of ULM4) could account for this difference by offsetting repulsion of the phosphorylated threonine. Despite the subtle penalty for affinity, phosphorylation enhanced the preference of U2AF2 for binding the SF3B1 ULM5 to 23-fold over the presumably off-target ULM4, which could facilitate specific U2AF2 interactions.

### Potential for switch-like regulation of U2AF2 by SF3B1 hyperphosphorylation

Processive phosphorylation of SF3B1, and the ensuing extreme negative electrostatic charge, could weaken U2AF2 association up to a critical point triggering its exit from the spliceosome (**Fig. 7**). Such all-or-none or switch-like regulation of IDRs by phosphorylation is well established.^44^ Temporally, SF3B1 phosphorylation levels are inversely correlated with detection of U2AF2 in the assembling spliceosome. SF3B1 is unphosphorylated in the A complex, moderately phosphorylated in the B complex, and then is highly phosphorylated in the B^act^ complex. At this stage, essentially all TP motifs in the SF3B1 ULM region are phosphorylated in preparation for the first catalytic step of pre-mRNA splicing.^3, 5–7^ Mass spectrometry (MS) analysis of catalytically activated B^act^ spliceosomes identified numerous, specific phosphorylation sites in the SF3B1 ULM region at this stage, including T211, T235, T313, T248, T326, and T328.^5^ In parallel, U2AF2 is required to initiate the E stage, detected in the A/B stages, and no longer evident in B^act^/C complexes.^3^ Furthermore, splicing extracts treated with a small molecule stabilizer of U2AF2 – PPT complexes accumulate H/E complexes, confirming that U2AF2 must be removed before spliceosome activation.^45^ Likely alongside other complementary regulatory functions, this evidence converges on hyperphosphorylation of the SF3B1 ULM region serving as a U2AF2-dependent gateway for pre-mRNA splicing to proceed.

The glutamate substitutions mimicking complete TP-phosphorylation in the SF3B1 ULM region abolished its detectable association with the U2AF2 protein. In contrast, omitting the second phosphorylation site in the 1(P)ULM4 peptide as compared to 2(P)ULM4 did not significantly change its binding affinity for the U2AF2 UHM. Of the two phosphorylated TP motifs present in the co-crystallized 2(P)ULM5 peptide, the C-terminal (P)TP motif was absent from the electron density. Yet, the dramatic penalty for U2AF2 binding following complete “phosphorylation” of the SF3B1 region is not due to disrupted protein folding since the ULM region is an IDR. Moreover, alanine substitutions for the phosphorylated threonines had no significant effect on SF3B1 binding to U2AF2.

Instead, global electrostatic repulsion is likely to play a role in the strong repulsive effect of complete phosphorylation or (P)-mimics in the SF3B1 ULM region on U2AF2 association in the context of the full length or nearly full length proteins. Phosphorylation of all 28 TP motifs converts the SF3B1 ULM region from a slight net positive charge (+3 unmodified) to a high negative charge (-47 phosphorylated) at physiological pH (7.4). Likewise, a large negative charge (-25) is expected for the engineered (P)-mimic SF3B1 variant, although less dramatic than the net charges from 28 doubly ionizable phosphoryl groups. The extreme negative charge following complete phosphorylation of the SF3B1 ULM would repel U2AF2, which is unusually acidic for an RNA binding protein, bearing negative charges of -10 and -11 for the UHM and U2AF^123L^ regions used for ITC. Indeed, a low isoelectric point is a feature that distinguishes UHMs from the similarly folded RNA recognition motif,^20^ suggesting that electrostatic repulsion by complete phosphorylation of the SF3B1 ULM region might apply to other members of the UHM family.

In line with the chromatin association of phosphorylated SF3B1, U2AF2, and the co-transcriptional splicing of most nascent pre-mRNAs,^5, 15, 46, 47^ the TP motifs in the SF3B1 ULM region are documented substrates of RNA polymerase II (RNAPII)-associated CDKs. CDK11 phosphorylates at least half of the TP motifs in the SF3B1 ULM region, including Thr211, Thr235, Thr313, and Thr328. These phosphorylation events are necessary for SF3B1 to associate with the U5 and U6 snRNAs for spliceosome assembly.^16^ CDK7 influences phosphorylation of 16 TP motifs in the SF3B1 ULM region, and inhibition of CDK7 alters pre-mRNA splicing.^8^ Moreover, treatments with CDK9 or CDK12/CDK13 inhibitors decrease SF3B1 phosphorylation levels and weaken its association with RNAPII.^10–12^ The sequential activities of CDK7, CDK9, CDK11, and CDK12/13 in the transcriptional stages of initiation, elongation, termination, and 3’ end processing highlight the importance of future studies to dissect the interplay of SF3B1 phosphorylation sites and the transcription-coupled recruitment of the U2 snRNP.

### SF3B1 (P)-variants disrupt pre-mRNA splicing

Our establishment of SF3B1 (P)-variants that either prevent or mimic TP-phosphorylation in the ULM region, which is the predominant phosphorylated domain of SF3B1, offer tools to investigate the transcriptome-wide effects of these phosphorylation states on pre-mRNA splicing. Expression of the (P)-variants as opposed to WT SF3B1 primarily led to cassette exon skipping or intron retention events that did not rescue, but indeed significantly overlapped, differential splicing events following endogenous SF3B1 depletion. By Occam’s Razor, the mechanistic action of SF3B1 phosphorylation releasing U2AF2 can explain the shared effects of SF3B1 depletion, (P)-dead, and (P)-variant SF3B1 (**Fig. 7**). First, we previously established significant overlap between splicing events affected by U2AF2 and SF3B1 depletions.^22^ Furthermore, chromatin-bound U2AF2 enhances exon inclusion in highly expressed genes,^46^ which is the converse of the numerous exon skipping events observed here. Second, (P)-dead SF3B1 is expected to stall spliceosome assembly at an early (E)-stage containing U2AF2, which cannot proceed to catalysis. Third, (P)-mimic SF3B1 might not sufficiently recruit U2AF2, which likewise would disrupt spliceosome assembly and pre-mRNA splicing at the major class of 3’ splice sites. These distinct, U2AF2-dependent mechanisms for splicing inhibition by (P)-dead or (P)-mimic SF3B1 are further supported by external evidence. For instance, treatments with either a compound that stabilizes the U2AF2 – PPT complex,^45^, a CDK inhibitor,^16^ or depletion of phosphatases,^19^ inhibits pre-mRNA splicing by blocking spliceosome assembly at distinct H/E, B, or C stages. Moreover, SF3B1-targeted inhibitors that stall spliceosome assembly at the A-stage also increase SF3B1 phosphorylation,^48, 49^ a finding which our results indicate would lead to secondary effects on association with U2AF2.

In agreement with the sensitivity of U2AF2 association to SF3B1 phosphorylation, the (P)-variant SF3B1 often penalized splicing of sites with uridine-rich PPTs, the preferred binding site of U2AF2.^30, 39, 40^ The PPT signals preceding cassette exons with increased skipping (i.e., decreased splicing of the proximal 3’ splice site) were significantly enriched in uridines following re-expression of (P)-dead or (P)-mimic SF3B1 (relative to WT SF3B1). Compared to the SF3B1-depleted samples, the PPTs of the retained introns with decreased splicing in the presence of the (P)-variant SF3B1 also were significantly enriched in uridines, possibly reflecting a greater dependence on U2AF2. Considering that polyuridine is an unstructured RNA sequence,^50, 51^ the SF3B1-specific retention of introns with uridine-poor PPTs could reflect loss of SF3B1-associated RNA helicase activities to chaperone structured, degenerate PPTs for the progression of spliceosome assembly (e.g., DDX42, hPrp5/DDX46, or hPrp43/DHX15)^52, 53^.

A small subset of sites exhibited increased splicing among the various SF3B1 samples. The PPTs with increased splicing in the presence of the (P)-variants, particularly (P)-dead SF3B1, lost the uridine-rich signature of U2AF2 regulation and instead were degenerate. Essentially all the differentially spliced sites were marked by U2-type splice site sequences rather than those of the minor U12-type spliceosomes. Notably, RNA secondary structure in 3’ splice sites can replace the need for U2AF2 during pre-mRNA splicing.^54^ Moreover, cancer-associated mutants of SF3B1, which typically affect residues distant from the U2AF2-interacting region and instead contact the PPT,^4^ increase use of 3’ splice sites in otherwise inaccessible RNA structures.^55^ Therefore, the rare subsets of degenerate PPTs for which the (P)-variants increased, rather than decreased splicing, are likely to be governed by kinetics and contexts involving other SF3B1 modulators in the major spliceosome. For example, U2AF1 is known to regulate 3’ splice sites with weak PPTs.^31^

### Implications of SF3B1 phosphorylation for its dysregulation in cancers

A strong relationship between SF3B1 and human disease raises the intriguing question of whether SF3B1 phosphorylation and/or its modulation of U2AF2 would be pragmatic targets for future therapies. Dysregulated pre-mRNA splicing is a long-recognized hallmark of cancers.^56^ SF3B1 is considered the most frequently mutated splicing factor gene across various cancer types, particularly myelodysplastic syndromes (MDS) with ring sideroblasts, chronic lymphocytic leukemias (CLL) and uveal melanomas.^57, 58^ Hotspot mutations located in the core, α-helical repeat region of the SF3B1 protein, particularly K700E, neo-functionally alter branch site choice,^59^ disrupt the E-to-A complex transition, dysregulate RNAPII kinetics,^60^ and increase pathological R-loops.^61, 62^

Despite the distinct locations of its cancer-associated mutations and major phosphorylation sites, SF3B1 phosphorylation is likely to both influence and be influenced by its dysregulated functions in cancer. Remarkably, a recent study noted elevated SF3B1 phosphorylation in CLL cells compared to normal B cells, irrespective of SF3B1 mutational status.^49^ Treatment with an SF3B1 inhibitor decreased phosphorylated SF3B1 levels without altering the total amount of SF3B1 protein. This study thereby connects the phosphorylation state of SF3B1, rather than its mutational status, with dysregulated pre-mRNA splicing in CLL and demonstrates its responsiveness to therapeutic modulation.

Our discovery that U2AF2 dissociation is sensitive to SF3B1 phosphorylation further suggests that compounds disrupting this interface or mimicking its phosphorylation could selectively eliminate “splicing-sick” cancer cells. Already, several breakthrough studies have demonstrated that specific inhibitors of transcriptionally-associated CDKs, such as SY-351 (for CDK7),^8^ OTS964 (for CDK11),^16^ or THZ531 (for CDK12/13),^12^ decrease SF3B1 phosphorylation and pre-mRNA splicing. Future work is needed to explore the full therapeutic potential for both U2AF2 modulation,^45^ and selective inhibitors of transcription-associated CDKs (e.g., KB-0742, SY-5609, or AZD4573 for CDK9; samuraciclib or enitociclib for CDK7) currently in clinical trials^63^ to target cancers with increased SF3B1 phosphorylation. The clinical success of CDK inhibitors as a drug class, exemplified by agents like palbociclib, ribociclib, and abemaciclib which target CDK4/6 and are widely used in breast cancer therapy, underscores the therapeutic potential of modulating protein phosphorylation to treat cancer.

In summary, we have shown that phosphorylation of the SF3B1 ULM region contributes to its release of U2AF2, thereby clearing the path for pre-mRNA splicing to proceed. This work establishes a new paradigm whereby protein phosphorylation, as a regulatory mechanism distinct from the ATP-dependent actions of RNA helicases, directly regulates key protein-protein interactions essential for spliceosome activation. Future studies will be needed to dissect the spatial and temporal regulation of SF3B1 phosphorylation within the full spectrum of cellular contexts and signaling pathways that control SF3B1 phosphorylation *in vivo*, particularly in diseased states. Ultimately, these findings will be instrumental in understanding spliceosome regulation by protein phosphorylation and could offer new avenues for therapeutic intervention in splicing-related diseases.

## METHODS

### Proteins and peptides

Recombinant proteins produced in this study were derived from NCBI RefSeq IDs NP_036565 (SF3B1) and NP_009210 (U2AF2). The U2AF2 UHM (residues 371-471), U2AF2^12UL^ (residues 141-471), and SF3B1 (residues 147-462) constructs were expressed and purified for crystallography and ITC as described previously.^22^ Full-length FLAG-tagged SF3B1 and HA-tagged U2AF2 were used for transient transfections as described previously.^22^ Variations were constructed and sequenced by GenScript Biotech Corporation. Twenty-eight threonines in TP motif clustered at the SF3B1 ULM region (**Fig. 1a**) were mutated either to alanines (phospho-dead SF3B1) or glutamates (phospho-mimic SF3B1). Phosphorylated synthetic peptides were synthesized and purified by Bio-Synthesis, Inc. The SF3B1 ULM5 peptide comprises residues 333-351, and 2(P)-ULM5 is phosphorylated on residues T341 and T350. The SF3B1 ULM4 peptide comprises residues 288-304, and 2(P)-ULM4 is phosphorylated on residues T296 and T303. The 1(P)-ULM4 peptide comprises residues 288-302, phosphorylated on residue T296. Peptide sequences are inset in **Fig. 1c**.

### Isothermal titration calorimetry

All proteins were dialyzed overnight at 4 °C in 4 L of buffer comprising 50 mM NaCl, 25 mM HEPES pH 7.4, 0.2 mM TCEP the day before ITC experiments, and peptides were diluted >50-fold into the dialysis buffers. ITC experiments used either a MicroCal VP-ITC or PEAQ-ITC (Malvern Panalytical). At least two replicates of the ITC experiments were run at 30 °C. The SF3B1 variant was used in the sample cell at 4-5 μM concentrations, except the phosphorylated peptides were used at 30-45 μM concentrations to account for the low binding affinities. The U2AF2 proteins were placed in the syringe at 10-fold higher concentrations. The “c-values” of the isotherms (c = n K_A_ [M], where n is the apparent stoichiometry, K_A_ is the apparent equilibrium binding affinity, and [M] is the concentration of macromolecule in the sample cell) were within reliable limits (9 ≤ c ≤ 100). Isotherms were corrected for heats of dilution by globally subtracting the average heats of the last 3-5 injections. After removing an initial test injection, the isotherms were fit using a single-site binding model using Origin® v7.0 implemented for the VP-ITC or the PEAQ-ITC software. The isotherms are shown in **Supplementary Fig. 1** and ITC results reported in **Supplementary Table 1**.

### Crystallization and structure determination

The U2AF2 UHM (20 mg mL^-1^) was mixed with 1.5-fold molar excess of SF3B1 peptide and crystallized by the hanging drop vapor diffusion method at 4 °C. For both structures, the reservoir solution contained 0.1 M sodium citrate tribasic dihydrate pH 5.6, 5% v/v 2-propanol, 30% v/v PEG 3350 and 0.2 M ammonium acetate. The crystals were cryoprotected by sequential transfer to 20% v/v glycerol in reservoir solution and maintained at 100 K for remote data collection using beamline 12-2 of the Stanford Synchrotron Radiation Light (SSRL) source. Data sets were processed using XDS^64^ and CCP4 packages^65^ implemented in the SSRL AUTOXDS script (A. Gonzalez and Y. Tsai). The structures were determined by molecular replacement using Phaser^66^ as implemented in Phenix^67^ and the U2AF2 UHM as the search model. Top solutions were refined using Phenix^67^ and manually adjusted in Coot.^68^ The crystallographic data collection and model quality statistics are reported in **Supplementary Table 2**.

### Cell culture, transfections, and co-immunoprecipitations

HEK 293T cells (ATCC^®^ CRL-3216^TM^) were maintained and transfected as described previously.^22^ For knockdown experiments, cells were harvested two days after transfection with Stealth^TM^ siRNAs (25 nM, Thermo Fisher Sci.) targeting SF3B1 (Cat. nos. HSS146415 is siSF3B1 in **Figs. 4 - 5** and siSF3B1a in **Supplementary Fig. 5 - 6**; HSS146414 is siSF3B1b in **Supplementary Fig. 5 - 6**) or “Low GC” control (Cat. No. 12935200). For “rescue” experiments with SF3B1 variants, the samples were harvested two days after co-transfection of the siRNAs and plasmid DNAs. Co-immunoprecipitation methods were variations of those previously described,^22^ with additional phosphatase inhibitors (0.1 mM sodium orthovanadate and 5 mM sodium fluoride as well as 50 mM β-glycerophosphate, MilliporeSigma) and washed with IP buffer containing MgCl_2_ and Turbonuclease (50 U, MilliporeSigma). For phosphatase experiments, cells were lysed in buffer without phosphatase inhibitors and split. Prior to co-immunoprecipitation, cell lysates were treated either with Lambda PPase (400U, NEB) in buffer containing 1 mM MnCl_2_ for 1hr, or phosphatase inhibitors were added, as indicated.

### Immunoblotting

For immunoblots of total protein (**Supplementary Figs. 3-5**), harvested cells were lysed and blotted as described^22^ with antibodies specific for SF3B1 (Abcam, Cat. No. ab170854 or MBL, Cat. No. D221-3), phosphoryl-TP (Cell Signaling Technology, Cat. No. 9391), phosphoryl-T211 SF3B1 (Invitrogen, Cat. No. PA5105427), phosphoryl-T313 SF3B1 (Cell Signaling Technology, Cat. No. 25009), U2AF2 (Sigma-Aldrich, Cat. No. U4758), FLAG (Sigma-Aldrich, Cat. No. F1804 or Invitrogen, Cat. No. 740001), HA (Sigma-Aldrich, Cat. No. H6908), H2B (Cell Signaling Technology, Cat. No. 12364), or GAPDH (Cell Signaling, Cat. No. 2118). Antibodies were diluted 1:1000 v/v with 5% w/v dry milk or BSA in TBS-T. Secondary antibodies included anti-rabbit IgG horseradish peroxidase (Invitrogen, Cat. No. 31460) or anti-mouse IgG horseradish peroxidase (Invitrogen, Cat. No. 31340). The chemiluminescence signal from Clarity^TM^ Western ECL substrate (Bio-Rad, Cat. No. 170-5061) was detected on a Chemidoc^TM^ MP Imaging System (Bio-Rad). Membranes were essentially cut as shown before blotting with different antibodies.

### RT-PCR of endogenous gene transcripts

The RT-PCR protocols and primer sequences were described previously^22^. Briefly, total RNA was isolated from transfected cells and DNase I-treated using an RNeasy Plus kit (Qiagen). The cDNAs were synthesized using random primers and Moloney murine leukemia virus RT (Invitrogen). Amounts of cDNA were normalized to 18S rRNA with the standard curve method, using quantitative real-time RT-PCR run with SYBER^TM^ Green in triplicate on a Bio-Rad CFX thermal cycler. The RT-PCR of *SAT1* was run for 40 cycles of 94°C for 30 s, 55°C for 10 s, and 72°C for 15 s; *THYN1* was run for 40 cycles of 94°C for 30 s, 57°C for 10 s, and 72°C for 15 s; RNF10 was run for 40 cycles of 94°C for 30 s, 55°C for 10 s, and 72°C for 15 s, all run on an Applied Biosystems 2720 thermal cycler (Thermo Fisher). The RT-PCR products were separated on a 2% w/v agarose-TBE gel and stained with ethidium bromide. Products were visualized using a Gel Doc XR+ gel documentation system (Bio-Rad). Band intensities of three technical replicates were quantified and background corrected using ImageJ^69^ software, representative of at least three biological replicates. Uncropped RT-PCR images are shown in **Supplementary Fig. 5b-d**.

### RNA library preparation and sequencing

Total RNAs were extracted from transfected cells in biological triplicates using an RNeasy Mini kit (Qiagen) and further purified with an RNA Clean & Concentrator kit (Zymo Research). Total RNA concentrations were determined using a NanoDrop One spectrophotometer, and RNA quality was assessed at the UR Genomics Research Center (URGRC) with an Agilent Fragment Analyzer. The RNA quality numbers were 10.0 for all samples. The URGRC prepared polyA-selected libraries using the TruSeq Stranded mRNA Sample Preparation Kit (Illumina) according to the manufacturer’s instructions. A NovaSeq 6000 high throughput sequencer (Illumina) was used to generate 150 bp paired-end reads for each sample. Raw reads were pre-processed at the URGRC with fastp^70^ 0.23.1 for adapter trimming and quality filtering. Reads were then aligned to the GRCh38 genome assembly using STAR^71^ 2.7.9a with basic two-pass mapping and removal of noncanonical intron motifs. Output BAM files were then sorted with Sambamba^72^ 0.6.8. Knockdown efficiencies for siSF3B1a and siSF3B1b were calculated using ratios of DESeq2^73^ normalized SF3B1 read counts (raw counts quantified by subread featureCounts^74^) averaged over replicates of siCtrl, siSF3B1a, and siSF3B1b samples.

### Bioinformatics and statistical analysis of RNA sequencing data

Protocols for identifying alternative splice events were variations of previously described methods.^75^ In short, splice junctions, identified by STAR,^71^ were filtered to include only those supported by a minimum of 15 uniquely mapped reads summed across the three replicates within each condition. These junctions were further filtered and assigned to genes if they mapped exclusively to single-gene loci as annotated by GENCODE Release 45.^76^ Splice event detection was implemented in a custom R script (version 4.2.1)^77^ running the tidyverse^78^, magrittr,^79^ and GenomicRanges^80^ packages. Across each condition pair of interest, junctions from both conditions were pooled and grouped into splicing events. Sets containing multiple junctions were then classified into one of the following alternative splicing types based on their junction configurations: cassette exon, alternative 5ʹ splice site, alternative 3ʹ splice site, or mutually-exclusive exons (MXE). Non-overlapping junctions were used to quantify intron retention. An additional filtering step ensured that each of the two conditions had detected all junctions necessary to complete at least one of the two outcomes for every splice event. Further analysis excluded MXE events, due to their low detection rate and infrequent differential significance, as well as complex events, which did not meet the criteria for these types. For input to rMATS, tables defining all identified alternative splice events were constructed from exon coordinates, which were inferred from positions of junctions in continuous regions of genes or for terminal junctions, GENCODE annotated exons. A custom GTF file was generated by appending newly detected splice event transcripts and exons to the GENCODE Release 45^76^ annotation. These “fixed event sets” and aligned reads (BAM files) were quantified and analyzed of differential splicing using rMATS-turbo 4.1.1.^81^

For **Fig. 5b** and **Supplementary Fig. 6a**, read ratios were calculated from the read counts of the inclusion and skipped cassette exon isoforms of *THYN1, SAT1* and RNF10, which were obtained from rMATS JCEC results without custom event detection. The *SAT1* and *RNF10* cassette exons overlap complex splicing patterns and so were excluded from further analysis by the filtering process described above. For the histograms and Euler diagram in **Fig. 5c-d**, splice events were considered shared across different condition pairs if the genomic coordinates of the target feature (e.g., cassette exon, region between proximal and distal alternative splice sites, or retained intron) and the inner edges of flanking exons were identical, and if they exhibited the same direction of the change in splicing (e.g., increased inclusion, decreased retention). To calculate the Spearman correlation coefficients corresponding to the Euler plots (**Fig. 5c**, **Supplementary Fig. 6b**), splice events were considered for pairwise correlation if they passed significance thresholds for at least one of the two condition pairs being assessed. Ranking was performed by sorting ΔPSI values and using False Discovery Rate (FDR) to break ties such that events were arranged from most significant positive ΔPSI events to most significant negative ΔPSI events.

Histograms of differential splice event counts (**Fig. 5d**) were created with the ggplot2^82^ R package. Euler diagrams depicting differential splicing overlap were generated using the eulerr^83^ R package from differential splice events with thresholds for statistical significance of FDR Σ0.05, |ΔPSI| ≥0.10 for one or more variants compared to the corresponding control (siSF3B1 *vs.* siCtrl; (P)-dead *vs.* WT; (P)-mimic *vs.* WT) and FDR Σ0.05, |ΔPSI| ≥0.05 for the others. The ΔPSI (Percent Spliced In) measure, shown on the x-axis of **Fig. 5d**, was calculated by rMATS for each splice event type. By convention, a positive ΔPSI for a cassette exon indicates higher inclusion (more splicing of the preceding junction) in the first sample compared to the second, whereas for intron retention, it indicates higher retention (less splicing) in the first sample compared to the second. Read coverage plots were generated using IGV^84^ by summing BAM files across all replicates.

Gene ontology enrichment of biological pathways (**Fig. 5e**) was analyzed using ShinyGo^85^ on gene lists comprising these differential splice events against a background of all detected genes, and the outputs were plotted using GraphPad Prism. Sequence logos (generated with WebLogo^86^ 3.7.8) along with uridine contents for a window (-2 to -20 nucleotides) before 3’ splice sites were derived from sequences extracted from the GRCh38 primary genome assembly (GENCODE Release 45^76^).

Box plots of uridine content in **Fig. 6** were created with the ggplot2^82^ R package. For statistical analysis of uridine contents, the intron retention events were additionally filtered for at least 5% average retention across test and background conditions prior to uridine content comparisons to exclude introns with minimal or no retention (only nonsignificant intron retention events affected due to |ΔPSI| ≥0.10 significance threshold).

### Statistical significance calculations

Unpaired t-tests with Welch’s correction were calculated using GraphPad Prism v10 and results were indicated with asterisks as defined for **Fig. 1c**, **Fig. 4**, **Fig. 5b**, and **Supplementary Fig. 6a**. FDR values were calculated by rMATS^81^ or ShinyGO.^85^ The Spearman rank correlations and Mann-Whitney U tests were calculated with the R stats package (‘cor()’ and ‘wilcox.test()’ functions, respectively).^77^

## Data Availability

The X-ray crystallography data and coordinates generated in this study have been deposited in the Protein Data Bank with accession codes pdb_00009p8j and pdb_00009p8k. The raw and processed RNA-seq data generated in this study have been deposited in the Gene Expression Omnibus (GEO) database under accession number GSE304018. The supplementary data, including the detailed table of ITC results, lists of differential splicing events, and custom R scripts used for the analysis, have been uploaded to a Figshare repository for public release with a Digital Object Identifier (DOI) upon publication.

## Code Availability

Custom R scripts for RNA-seq data analysis have been provided as supplementary information for review and uploaded to a Figshare repository for public release with a Digital Object Identifier (DOI) upon publication.

## Acknowledgments

This study was supported by NIH R01 GM070503 and a Wilmot Cancer Institute GEM Pilot award to C.L. Kielkopf and NIH R01 GM141544 to P.L.B. H.R.P. and J.W.G. were supported by NIH T32 GM135134 and an Elon Huntington Hooker Fellowship. Crystallographic data were collected remotely using the Stanford Synchrotron Research Laboratory, supported by US DOE (Contract No. DE-AC02–76SF00515) and NIH (P41 GM103393).

## Author contributions

C.L. Kirchhoff accomplished RNA-seq analysis with guidance from P.L.B. H.R.P. accomplished the RT-PCR experiments and AlphaFold modeling. M.J.P. prepared RNA-seq samples, performed co-IPs, and directed RT-PCR experiments. S.L. and J.W.G. accomplished ITC, crystallization, and structure determinations. J.L.J. directed ITC and structure determinations. C.L. Kielkopf cryo-protected crystals, collected crystallographic data, and analyzed structures. C.L. Kielkopf conceived the study, designed the experiments, and wrote the manuscript with constructive input from all authors. The authors confirm adherence to ethical standards in the conduct of this research, including transparent reporting of all data and proper attribution of contributions.

