## Supplementary for "SF3B1 Phosphorylation Prompts U2AF2 Dissociation for Widespread Control of pre-mRNA Splicing"

**Contents:**

Supplementary Table 1. Isothermal titration calorimetry results
Supplementary Table 2. Crystallographic data and refinement statistics
Supplementary Table 3. Number and significance of differential splicing events

Supplementary Fig. 1. Representative isotherms of ITC experiments
Supplementary Fig. 2. AlphaFold predictions of the SF3B1 ULM region and its variants.
Supplementary Fig. 3. Immunoblots for phosphorylation levels of SF3B1 variants
Supplementary Fig. 4. Co-immunoprecipitations of (P)mimic SF3B1 and SPF45 or Tat-SF1
Supplementary Fig. 5. Immunoblots and uncropped gel images of RT-PCR samples
Supplementary Fig. 6. Immunoblots and comparison of siRNAs for RNAseq samples

Available to download as separate files:

Supplementary Data 1. List of differential splicing events for siSF3B1a *vs.* siCtrl
Supplementary Data 2. List of differential splicing events for siSF3B1b *vs.* siCtrl
Supplementary Data 3. List of differential splicing events for WT SF3B1 *vs.* siCtrl
Supplementary Data 4. List of differential splicing events for (P)-dead SF3B1 *vs.* WT SF3B1
Supplementary Data 5. List of differential splicing events for (P)-mimic SF3B1 *vs.* WT SF3B1

**Supplementary Table 1.** Isothermal titration calorimetry of U2AF2 regions titrated into SF3B1 regions.

| **n** | **$K_D$** (nM) | **ΔG** (kcal $mol^{-1}$) | **ΔH** (kcal $mol^{-1}$) | **-TΔS** (kcal $mol^{-1}$) |
|---|---|---|---|---|
| **$U2AF2^{12UL}$ titrated into wild-type SF3B1 ULM region:** [1] | | | | |
| 2.0 ± 0.1 | 800 ± 80 | -8.5 ± 0.1 | -9.4 ± 0.2 | 0.9 ± 0.3 |
| **$U2AF2^{12UL}$ titrated into phospho-dead SF3B1 ULM region:** | | | | |
| 4.1 ± 0.3 | 1,160 ± 134 | -8.2 ± 0.1 | -4.3 ± 0.1 | -4.0 ± 0.1 |
| **$U2AF2^{12UL}$ titrated into phospho-mimic SF3B1 ULM region:** | | | | |
| *No heats of binding detected* | | | | |
| **U2AF2 UHM titrated into SF3B1 ULM5:** [1] | | | | |
| 0.94 ± 0.1 | 46 ± 3 | -10.2 ± 0.0 | -18.4 ± 1.3 | 8.2 ± 1.4 |
| **U2AF2 UHM titrated into SF3B1 2(P)ULM5:** | | | | |
| 0.85 ± 0.0 | 190 ± 3 | -9.3 ± 0.0 | -18.4 ± 0.3 | 8.4 ± 1.3 |
| **U2AF2 UHM titrated into SF3B1 ULM4:** | | | | |
| 0.90 ± 0.0 | 680 ± 99 | -8.6 ± 0.1 | -13.8 ± 1.3 | 5.2 ± 1.4 |
| **U2AF2 UHM titrated into SF3B1 2(P)ULM4:** | | | | |
| 0.93 ± 0.0 | 4,360 ± 23 | -7.4 ± 0.0 | -12.9 ± 0.5 | 5.5 ± 0.6 |
| **U2AF2 UHM titrated into SF3B1 1(P)ULM4:** | | | | |
| 0.96 ± 0.0 | 3,720 ± 23 | -7.5 ± 0.1 | -13.0 ± 0.7 | 5.4 ± 0.8 |

Constructs boundaries and modifications are described in the methods and shown in Fig. 1.
Average values and standard deviations of 2-5 separate titrations are reported. Values for ULM5 include two measurements reported in Galardi *et al* (2022) *J. Biol. Chem.* v298:102224 plus three replicates for this work. All ITC experiments used the same buffer composition (50 mM NaCl, 25 mM HEPES pH 7.4, 0.2 mM TCEP).
n, apparent stoichiometries of the fit.
ΔG is calculated using the equation $\Delta G = -RT \ln (K_A)$ where T=303 K and $K_A$ is $1/K_D$.
-TΔS is calculated using the equation $-T\Delta S = \Delta G - \Delta H$.

**Supplementary Table 2. X-ray data collection and refinement statistics.**

| U2AF2 UHM with: | SF3B1 2(P)ULM5 | SF3B1 1(P)ULM4 |
|---|---|---|
| PDB Code: | 9P8K | 9P8J |
| **Data collection** | | |
| Space group | P $2_1$ | P 1 |
| Unit cell a, b, c (Å) | 58.5, 29.4, 58.7 | 48.7, 49.7, 60.8 |
| α, β, γ (°) | 90.0, 90.3, 90.0 | 74.8, 86.5, 65.5 |
| Resolution range (Å) (highest shell) | 29.44 – 1.43 (1.46 – 1.43) | 36.88– 2.30 (2.38 – 2.30) |
| Wilson B (Å$^2$) | 14.1 | 38.3 |
| $R_{measure}$ (%) | 9.2 (35.9) | 5.7 (24.3) |
| $CC_{1/2}$ (%) | 99.5 (92.3) | 99.6 (94.7) |
| Mean I/σ(I) | 11.0 (3.2) | 9.3 (3.3) |
| Completeness (%) | 95.0 (49.9) | 85.5 (81.7) |
| Multiplicity | 5.0 (3.2) | 1.8 (1.7) |
| **Refinement** | | |
| $R_{work}/R_{free}$ (%) | 18.3 / 21.6 | 19.0 / 24.2 |
| Coordinate error (Å) | 0.16 | 0.26 |
| No. Reflections (work/free) | 35,483/3,862 | 19,005/1,140 |
| Complexes per ASU | 2 | 4 |
| No. Non-hydrogen atoms | | |
| All atoms | 2098 | 3751 |
| Protein, U2AF2 | 1728 | 3318 |
| Protein, SF3B1 | 190 | 300 |
| Waters | 176 | 131 |
| Other solvent molecules | 4 | 2 |
| <B-factor> (Å$^2$) | | |
| All atoms | 22.2 | 52.8 |
| Protein, U2AF2 | 21.4 | 51.0 |
| Protein, SF3B1 | 31.4 | 74.2 |
| Waters | 30.9 | 50.0 |
| Other solvent molecules | 39.8 | 66.4 |
| R.m.s. Bonds (Å) | 0.007 | 0.002 |
| R.m.s. Angles (°) | 0.84 | 0.47 |
| Ramachandran (%) favored/allowed/outliers | 100.0/0.0/0.0 | 98.8/1.2/0.0 |
| Clash score | 4.2 | 2.4 |

Values in parentheses refer to the highest resolution shell. $R_{measure} = \sum_h \sum_l \{[(n/(n-1)]^{1/2} |I_{hl} - \langle I_h \rangle|\}/\sum_h \sum_l \langle I_h \rangle$ is the multiplicity-weighted R-factor, where n is the number of observations of the intensity I.
$CC_{1/2}$, correlation coefficient between intensities of random half-dataset.
$R_{work} = \Sigma_{hkl}||F_{obs}(hkl)|-|F_{calc}(hkl)||)/\Sigma_{hkl}|F_{obs}(hkl)|$ for the working set of reflections.
For $R_{free}$, either 10% of the data for (P)ULM5 or 6% of the data for (P)ULM4, were selected randomly for exclusion from all stages of refinement. No cutoff was applied on the data used for refinement.
Coordinate error was estimated in Phenix by the Maximum Likelihood method.
Clash score is the number of unfavorable atom-atom overlaps, ≥ 0.4Å per thousand atoms, calculated using the program MolProbity as implemented within the Phenix package.
"Other solvent molecules" includes an acetate ion for 2(P)ULM5 and sodium ions for 1(P)ULM4.

**Supplementary Table 3. Number of differential splicing events and significance of uridine contents in PPTs.** The number of splicing events (sample sizes) in each class is given, using FDR ≤0.05 and |ΔPSI| ≥ 0.10 to be considered a significant splicing change. The *p*-values were calculated by Mann-Whitney U tests.

| Cassette Exons (Upstream 3´ Splice Site) | | | | |
|---|---|---|---|---|
| Sample 1: **siSF3B1** (vs siCtrl) | Sample 2: (P)-dead (vs WT) | **Splicing Change** | ***p*-value** | **Significance†** |
| 3,390 events | 1,628 events | **Decreased** | **4.69e-06** | ***** |
| 3,191 events | 3,686 events | **n.s.** | 0.026 | * |
| 83 events | 73 events | **Increased** | **1.21e-06** | ***** |
| **siSF3B1 (vs siCtrl)** | **(P)-mimic (vs WT)** | **Splicing Change** | ***p*-value** | **Significance†** |
| 3,390 events | 1,680 events | **Decreased** | **5.46e-04** | *** |
| 3,191 events | 3,639 events | **n.s.** | 0.034 | * |
| 83 events | 54 events | **Increased** | **6.87e-03** | ** |
| **Retained introns** | | | | |
| Sample 1: **siSF3B1 (vs siCtrl)** | Sample 2: **(P)-dead (vs WT)** | **Splicing Change** | ***p*-value** | **Significance†** |
| 2,592 events | 3,076 events | **Decreased** | **1.70e-64** | ***** |
| 7,925 events | 9,443 events | **n.s.** | 0.51 | n.s. |
| 110 events | 83 events | **Increased** | **3.91e-05** | **** |
| **siSF3B1 (vs siCtrl)** | **(P)-mimic (vs WT)** | **Splicing Change** | ***p*-value** | **Significance†** |
| 2,592 events | 1,950 events | **Decreased** | **3.13e-58** | ***** |
| 7,925 events | 9,162 events | **n.s.** | 0.077 | n.s. |
| 110 events | 95 events | **Increased** | **9.75e-04** | *** |

† Significance of difference between Sample 1 and Sample 2:
n.s. ≥ 0.05; * <0.05; ** <0.01; *** <1e-3; **** <1e-4; ***** <1e-5

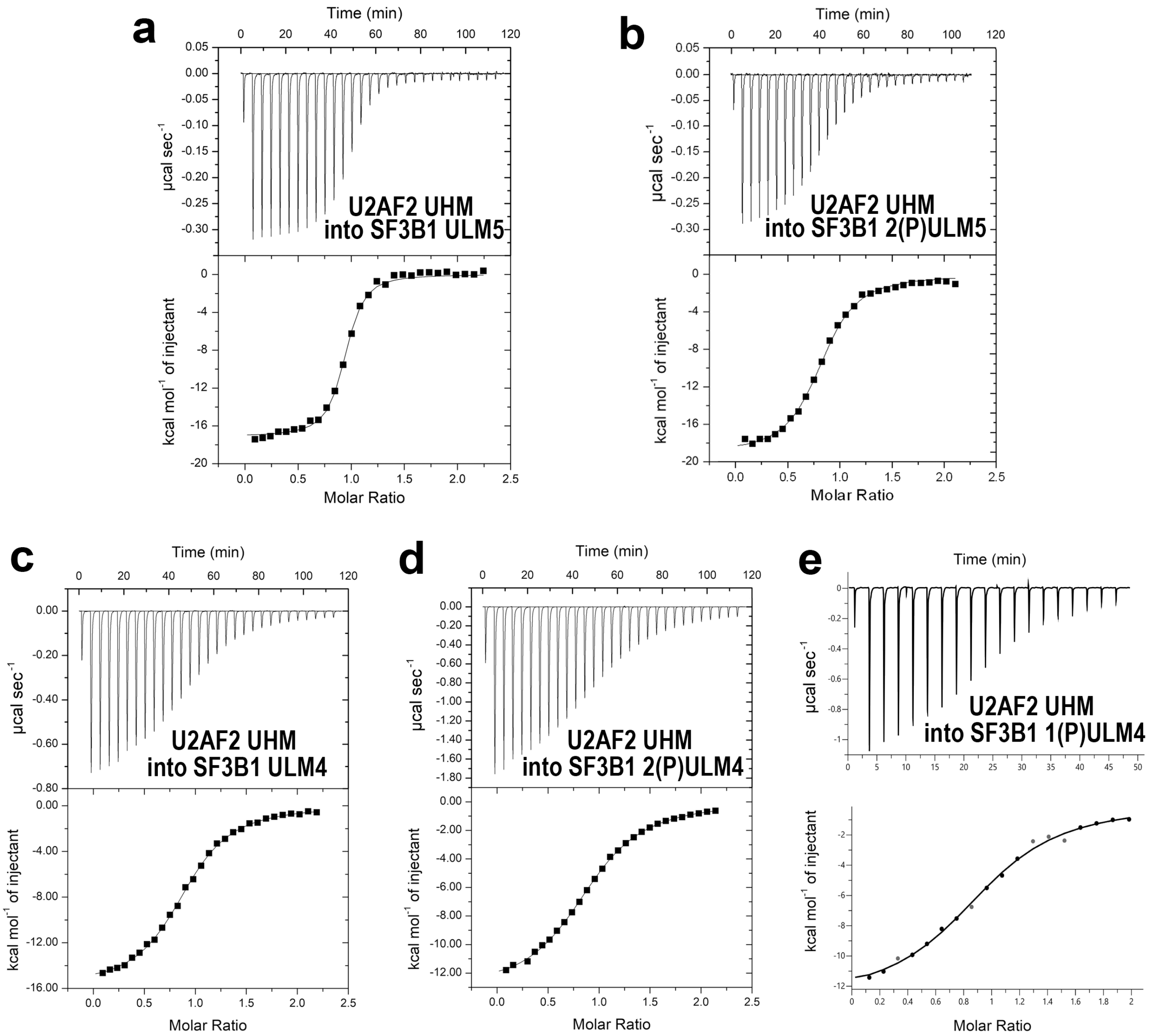


**Supplementary Fig. 1.** Representative isotherms of isothermal titration calorimetry experiments reported in **Fig. 1** and **Supplementary Table 1**. The U2AF2 UHM was titrated into **a** ULM5, **b** 2(P)ULM5, **c** ULM4, **d** 2(P)ULM4, or **e** 1(P)ULM4.

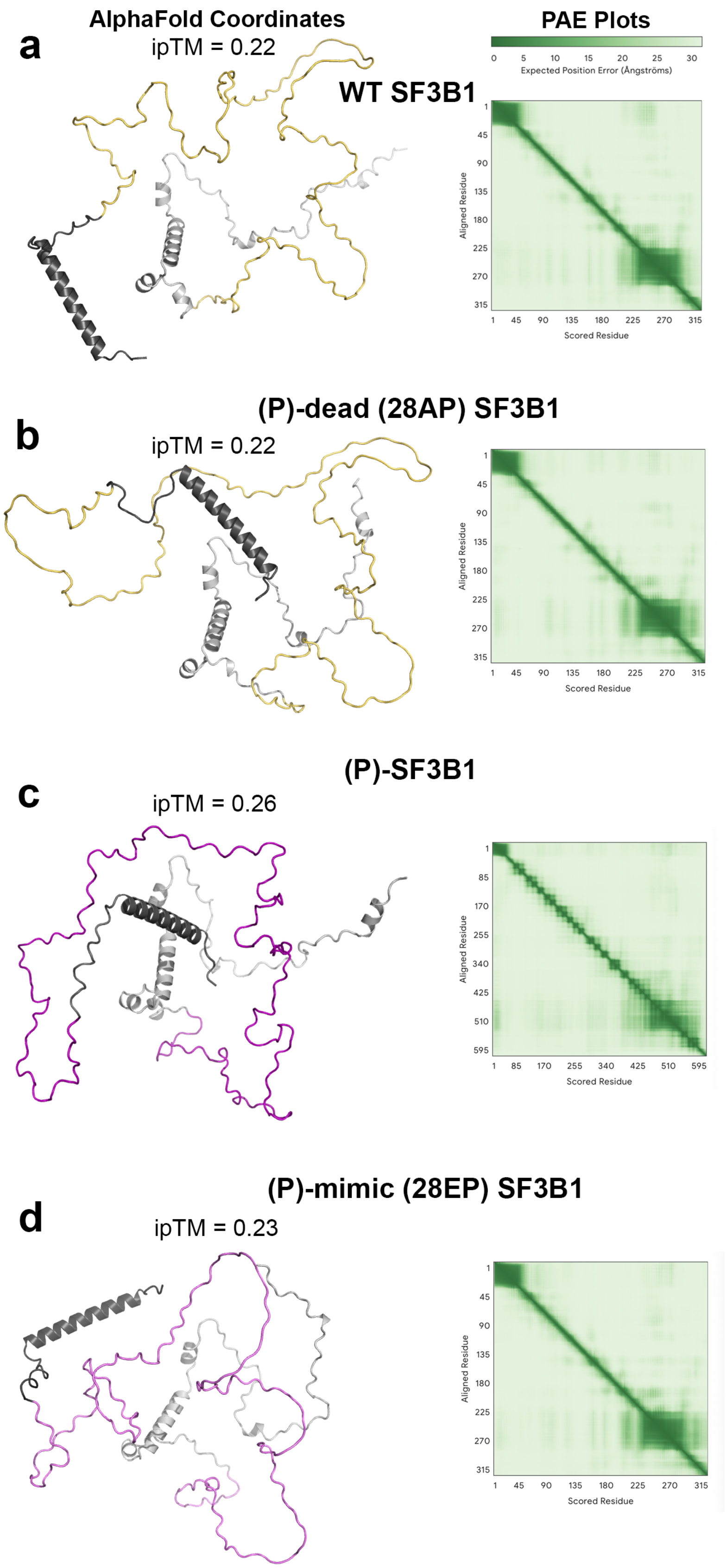

a
AlphaFold Coordinates
ipTM = 0.22
PAE Plots
0 5 10 15 20 25 30
Expected Position Error (Ångströms)
WT SF3B1
Aligned Residue
Scored Residue
b
(P)-dead (28AP) SF3B1
ipTM = 0.22
c
(P)-SF3B1
ipTM = 0.26
d
(P)-mimic (28EP) SF3B1
ipTM = 0.23

**Supplementary Fig. 2**. AlphaFold predictions of the recombinantly expressed SF3B1 ULM region (residues 147-462) and its variants, **a** WT, **b** (P)-dead, **c** phosphorylated, **d** (P)-mimic. The top ranked models are shown on the left with the interface predicted Template Modeling scores (ipTM) given above. The very low ipTMs reflect AlphaFold's lack of confidence in any fixed regular structure for this IDR. The Predicted Aligned Error (PAE) heat maps are shown to the right of each model. The very light color for residues in the ULM region (residues 47-211 of the model or 193-354 of full length SF3B1) reflect AlphaFold's lack of confidence in the placement of the ULM-containing part of the model relative to other parts. Residues 147-193 (dark gray) and 354-462 (light gray) flanking the ULM region were included in the construct to improve the stability of the recombinant protein. Residues 147-193 do not contain TP phosphorylation sites. Residues 354-462 contain $\alpha$-helical structures when bound to the SF3B6 subunit (PDB ID 2F9D), which are reflected in the structure prediction. The three phosphorylated TP motifs in this SF3B1 range fall in coil regions outside of the predicted $\alpha$-helices and the SF3B6 interface. Neither phosphorylation nor TP>AP/EP substitutions have any noticeable effect on the predicted secondary structures.

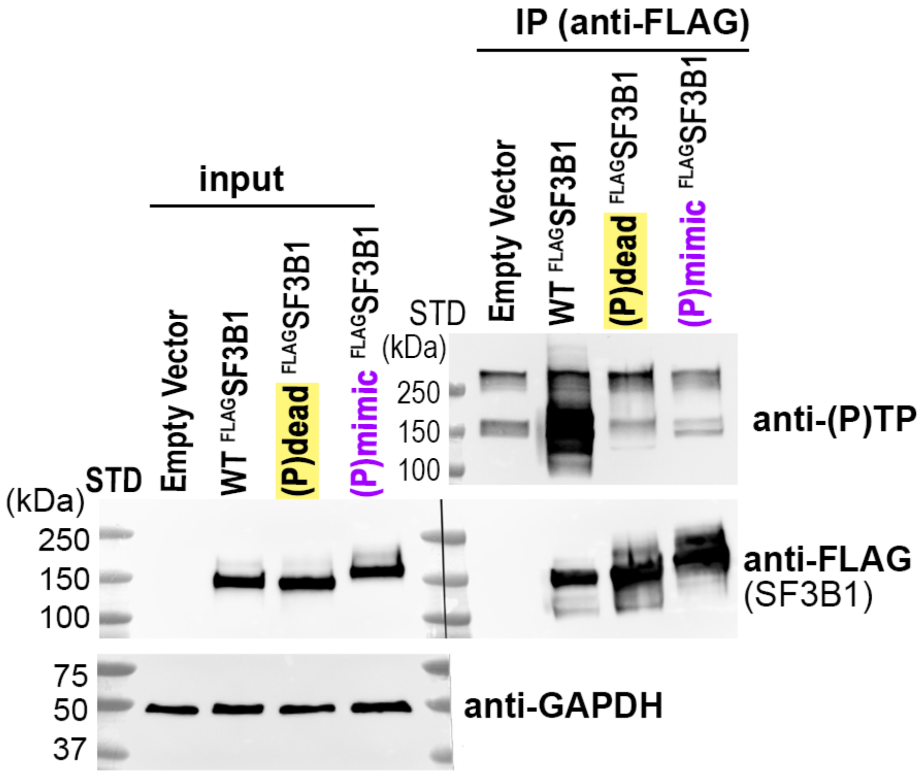


**Supplementary Fig. 3.** Immunoblots detecting phosphoryl-threonines or controls (FLAG, SF3B1, or GAPDH) of FLAG-immunoprecipitated samples expressing wild-type (WT), phospho (P)-dead, or (P)-mimic variants of $^{FLAG}$SF3B1 show that mutations in the ULM-region significantly reduce SF3B1 TP-phosphorylation.

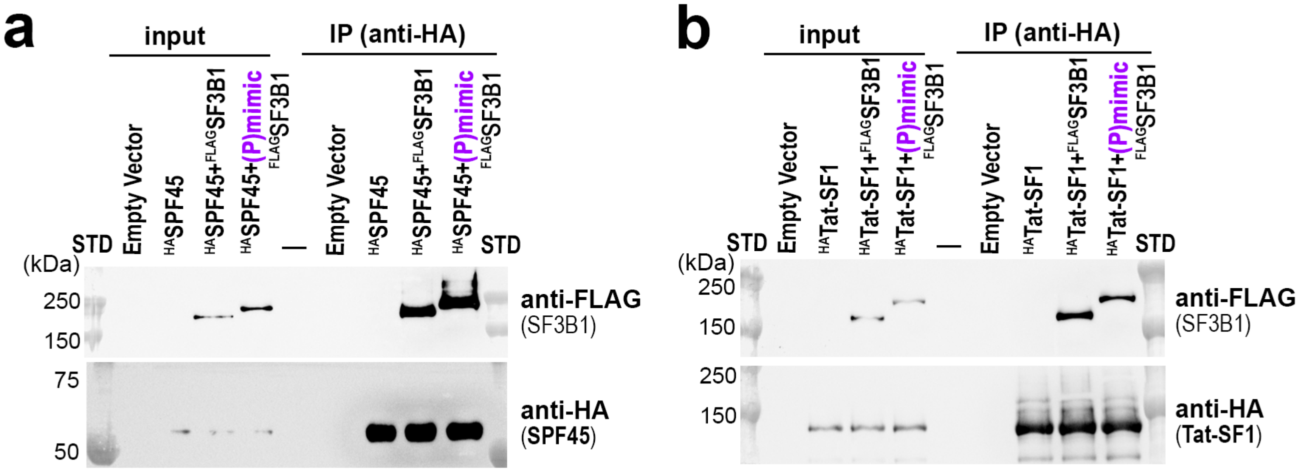


**Supplementary Fig. 4.** Co-immunoprecipitations of (P)-mimic Flag-tagged SF3B1 with either HA-tagged **a** SPF45 or **b** Tat-SF1 transiently expressed in HEK 293T cells. STD, protein molecular weight markers.

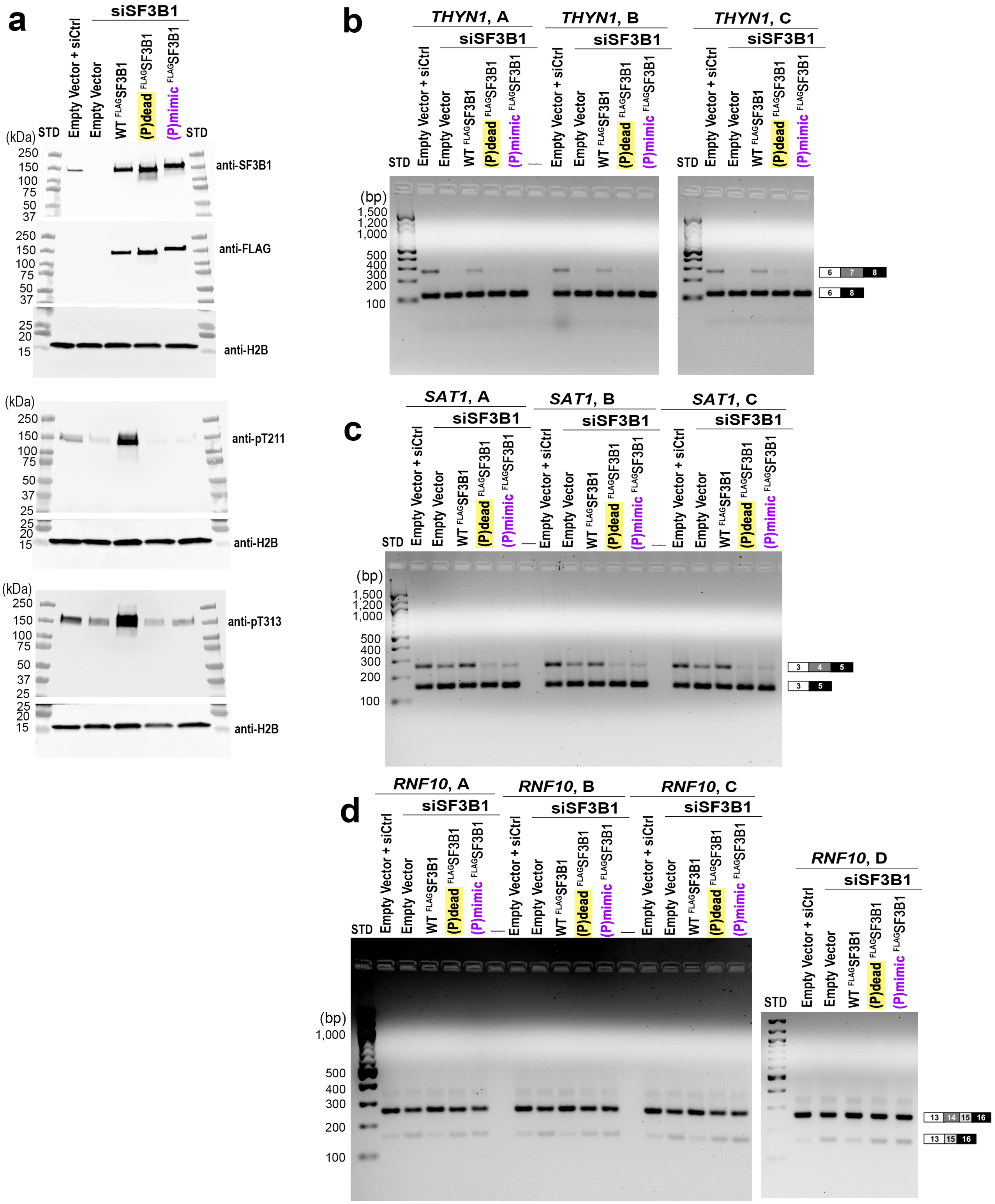


**Supplementary Fig. 5.** Supplementary data for the RT-PCR shown in Fig. 4. **a** Immunoblots of RT-PCR samples. (**b** – **d**) Uncropped gel images of RT-PCR replicates (labeled A, B, C) for **b** *THYN1,* **c** *SAT1,* **d** *RNF10*.

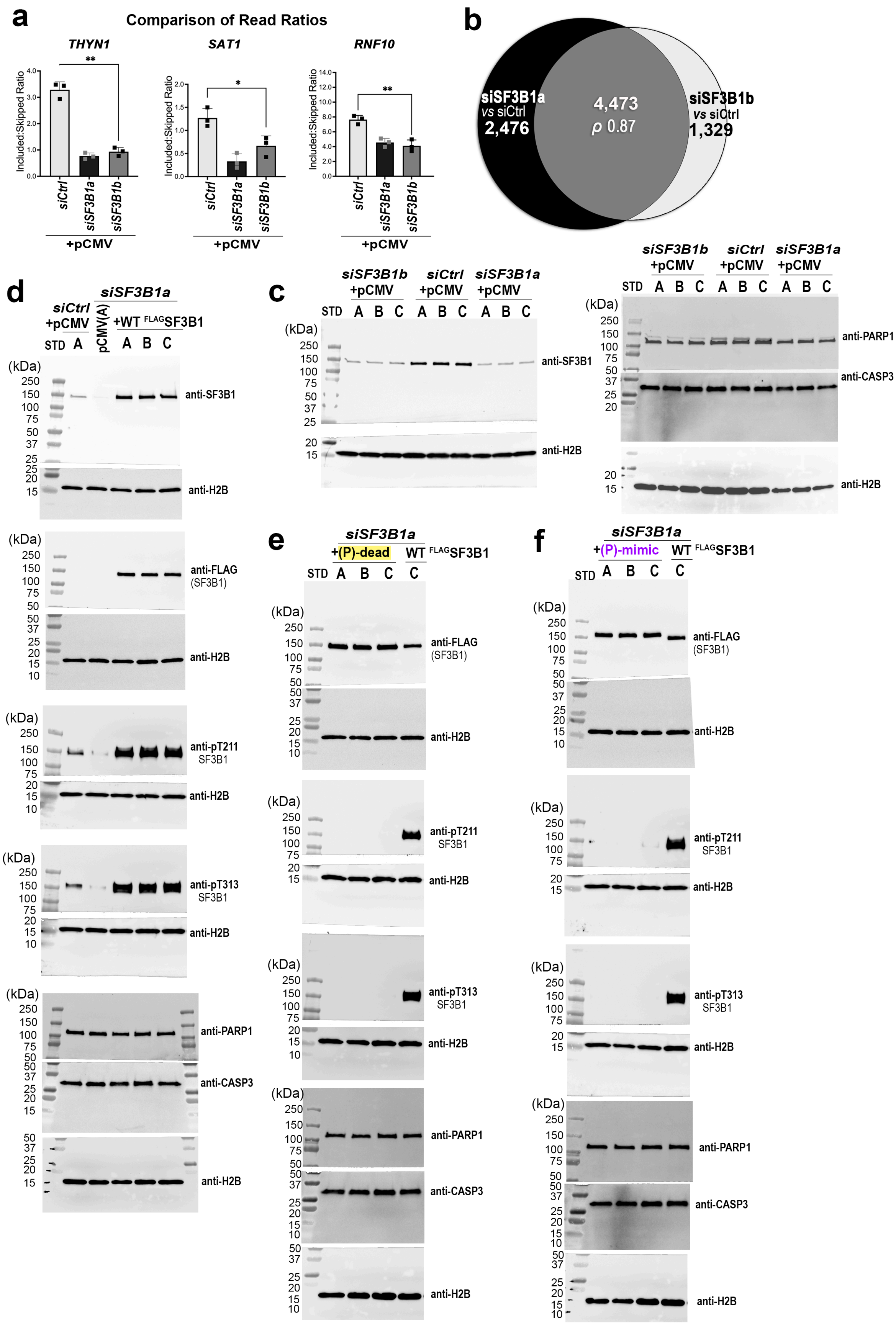

a
Comparison of Read Ratios
THYN1
SAT1
RNF10
Included:Skipped Ratio
siCtrl
siSF3B1a
siSF3B1b
+pCMV
b
siSF3B1a
vs siCtrl
2,476
4,473
ρ 0.87
siSF3B1b
vs siCtrl
1,329
c
siSF3B1b +pCMV
siCtrl +pCMV
siSF3B1a +pCMV
STD
A B C
(kDa)
anti-SF3B1
anti-H2B
anti-PARP1
anti-CASP3
d
siCtrl +pCMV
siSF3B1a
pCMV(A)
+WT FLAGSF3B1
anti-FLAG (SF3B1)
anti-pT211 SF3B1
anti-pT313 SF3B1
e
+(P)-dead
WT FLAGSF3B1
f
+(P)-mimic

**Supplementary Fig. 6.** Supplementary data for RNAseq analyzed in **Figs. 5** and **6**: **a** Quantified read counts for *SAT1*, *THYN1*, and *RNF10* for SF3B1 knockdown samples (*siSF3B1a*, black or *siSF3B1b*, light gray) compared to a control siRNA (*siCtrl*, unfilled) show similar SF3B1-sensitive changes as RT-PCR results. The empty vector (pCMV) for SF3B1-encoding plasmids is included for comparison with SF3B1-encoding plasmids. Welch's t-test: n.s., not significant; *, $p \leq 0.05$; **, $p \leq 0.01$. **b** Transcriptome-wide splicing changes are strongly correlated between the two SF3B1-targeted siRNAs. Cutoffs for statistical significance were FDR ≤ 0.05, $|\Delta PSI| \geq 0.10$, and the direction of the change in splicing (increased or decreased) was required to be the same. The Spearman's correlation coefficient (ρ) is inset. (**c – f**), Immunoblots of FLAG-tagged SF3B1, phosphoryl-T211 or phosphoryl-T313 of SF3B1, H2B loading controls, and confirming cell viability, PARP1 and CASP3, in samples used for RNAseq: **c** SF3B1 knockdown with re-expression of **d** wild-type (WT), **e** phospho (P)-dead, or **f** (P)-mimic SF3B1.
